# Identification of High Blanchability Donors, Candidate genes and Markers in Groundnut

**DOI:** 10.1101/2025.06.18.660288

**Authors:** Priya Shah, Sunil S. Gangurde, Ramachandran Senthil, Kuldeep Singh, Prashant Singam, Pasupuleti Janila, Sean Mayes, Manish K. Pandey

## Abstract

Blanchability in groundnut, the ability of seeds to shed their seed coat (testa), is a trait of economic importance in the food processing industry, yet remains underexplored in breeding programs. This study aimed to assess blanchability in 184 accessions from the ICRISAT minicore collection and identify associated genomic regions, candidate genes, and molecular markers. Significant variability was observed over two seasons, with blanchability ranging from 3.98% to 70.08%. Ten genotypes, including ICG10890, ICG9507, ICG13982, and ICG297, exhibited good blanchability, with ICG297 emerging as a promising donor based on cluster analysis of blanchability and agronomic traits. Genome-wide association studies (GWAS) using the 58K ‘Axiom_*Arachis*’ SNP array revealed 58 significant SNP-trait associations and important candidate genes such as *isocitrate dehydrogenase* and *ubiquitin ligase*, which influence seed coat structure and cell wall integrity. Nine SNPs were validated via allele mining, with four markers—on chromosomes A01 (snpAH00551), A06 (snpAH00554), B04 (snpAH00558), and B07 (snpAH00559), effectively distinguishing between high and low blanchability genotypes. These validated SNPs present valuable tools for genomics-assisted breeding. Overall, the findings is the first study contributing to a better understanding of the genomics and genetic basis of blanchability and offer resources for improving processing traits in groundnut.

## 1. Introduction

Groundnut (*Arachis hypogaea* L.) also known as peanut, is a major leguminous crop, grown mostly as a food and feed crop globally [1]. The cultivated species of groundnut is a self-pollinated allotetraploid crop (2n = 4x = 40) with 2.54 Gb genome size [2–4] and belongs to the *Fabaceae* family [5, 6]. It is a key seasonal herbaceous legume crop grown in arid and semi-arid regions and is recognized as one of the important sources of edible oil and protein. Groundnut kernels contain 16% to 36% of protein, 36% to 54% oil and between 10% and 20% of total carbohydrates [7]. Groundnut has significant market value and enormous nutritional value by virtue of high mono-unsaturated fatty acid (MUFA) in oil content and possess significant amount of vitamins and minerals [8]. It also possesses several nutritional qualities such as soluble sugars and possesses rich amounts of calcium (Ca), iron (Fe), Vitamin B and E (tocopherol). Additionally, anti-nutrients like trypsin inhibitor and phytic acid are present in groundnut that may be deactivated by the process of boiling and roasting [9, 10]. It has secured 13^th^ position among the most vital food crops and secured rank 4^th^ in the list of the most vital oilseed crops[11].

According to industrial importance, edible food products such as raw or roasted nuts, refined oil, groundnut butter, salted groundnuts, groundnut flour and other confectionery products are processed groundnut products. During the processing of groundnut to prepare the edible products, an important operation is the separation of the testa or seed coat (skin) and the process is called blanching. The blanchability is the capacity of a genotype to release its testa very easily. This characteristic is of great economic value in the processing of groundnut raw and processed food products since it reduces the cost of food processing. Additionally, this trait also holds importance in terms of quality control as blanchability leads to removal of damaged kernels, including aflatoxin contamination. Moreover, in terms of processing applications, globally, about 48% of groundnuts are used for the preparation of raw and processed food products such as groundnut milk, groundnut butter, groundnut chikki, groundnut cake, salted and roasted groundnut, groundnut bar chocolate, chutney, while oil extraction accounts for 52% usage, however in India, the groundnut utility accounts for food, seed, and oil extraction is 44%, 24% and 30% respectively [12, 13].

Blanchability has enormous economic significance in the production of food products made from groundnut. If the groundnut variety has low blanchability, the processing of the product becomes laborious, as it results in high re-processing and thus higher costs and time to get marketable products. This trait is associated with the seed coat and its structural properties as well as its chemical composition, such as lignin, pectin, polyphenols. Blanchability is reported to be a trait with high heritability and genetically regulated, hence the breeding and selection using conventional breeding could help in improving the groundnut cultivars with optimum blanchability (∼75%). In this context, GAB holds higher ability for accelerated improvement of crop varieties as the conventional plant breeding is prolonged and laborious [14]. Though there are several studies reported on germplasm screening for blanchability [14, 15], despite that only one study has reported the genomic regions for blanchability using QTL-seq approach on a RIL population [16]

SNP-trait association mapping of groundnut diverse panels has speed up the genomic region identification that are linked to agronomic traits by identifying ancestral or natural recombination events that led to non-random allele association at various loci throughout the genome [17, 18]. Compared to biparental linkage mapping, association mapping, respectively, allows greater mapping resolution [19]. With the availability of high-density SNP (Single Nucleotide Polymorphism) array and WGRS genotyping data, there is much scope to study the economically important and other traits in groundnut, identify STAs using the sequencing based molecular markers, and deploying them in crop breeding programs

Association mapping has acquired considerable pace in legumes, and there are several studies indicating markers associated with nutritional and agronomic traits [20]. Considering the above knowledge gap and available information, the present study aimed to identify STAs for economically important trait, blanchability, in the groundnut minicore set [21].

In this association mapping study, we have used three different algorithms, viz., Bayesian-information and linkage-disequilibrium iteratively nested keyway (BLINK), Settlement of Unmappable Positions via Extensive Rearrangement (SUPER), and Fixed and random model Circulating Probability Unification (FarmCPU). All these algorithms provide better statistical accuracy and computational efficiency and improve the accuracy of significant marker selection to be used in breeding programs.

Genome-wide association study (GWAS) has a potential to elucidate the genetic structure and the causative loci between different genotypes [22, 23]. This method has been widely utilized for the genetic dissection of complex traits in plants. As next-generation sequencing (NGS) technology has developed rapidly, high-density SNP can be invoked and additional GWAS analysis can be conducted. In the present study, a minicore collection comprising 184 diverse groundnut accessions were phenotyped for blanchability in two subsequent seasons (rainy (R) 2022 and post-rainy (PR) 2022-2023). We conducted GWAS for the identification of significant SNP markers related to blanchability in groundnut. Further, genes associated with blanchability were identified from the LD region of identified STAs. Also, the correlation and cluster analysis of blanchability with agronomic traits have revealed genotypes with superior traits. These results would provide a better and precise understanding of the blanchability trait with potential governing regions and its underlying molecular mechanism. Moreover, this study will further contribute to the improvisation of the blanchability of groundnut varieties with the help of marker-assisted breeding.

## 2. Material and Methods

### 2.1. Plant material and Phenotyping for blanchability

The groundnut minicore collection of 184 accessions developed at ICRISAT, Patancheru, was used in the experimental study [21]. Groundnut minicore collection encompasses six botanical types *viz*., *hypogaea*, *hirsuta*, *fastigiata*, *peruviana*, *aequatoriana* and *vulgaris,* that collectively constitutes the genetic diversity present in the complete groundnut germplasm at ICRISAT **[Supplementary Figure S4 (A)].** The experimental design used here was the alpha-lattice design for a total of two seasons: rainy (R) season of 2022 and post-rainy (PR) 2022-2023. Phenotyping data generated in two replications during each season at ICRISAT, Patancheru, (17° 31’ 48.00” N latitude and 78° 16’ 12.00” E longitude).

Each accession was planted in a 4 m row plot with inter-plant spacing of 10 cm and between-row spacing of 60 cm. The experimental site consists of red soils with pH ranging from 6.0 to 6.3. In the rainy season of 2022, the highest temperature was upto 31.26°C whereas the range of minimum temperature was recorded to be 20.38°C, with 198.2 mm rainfall, 154.24 mm of evaporation, and 87.60% accounts for relative humidity. But in the post-rainy season of 2022–2023, the highest temperature was 33.25°C, whereas the lowest temperature was recorded to be 18.61°C, with 65.5 mm rainfall, 98.75 mm rate of evaporation, and 82.70% of relative humidity. Throughout both the growth seasons, standard recommended agronomical practices essential for groundnut cultivation were followed for healthy and good crop growth. Further, post-harvesting, the phenotypic observations for blanchability were recorded for each replication at ICRISAT, Hyderabad, India **(Supplementary Table S1).** Additionally, the previously documented phenotyping data for various agronomical traits such as pods per plant (PPP), seed length (SDL) seed weight (SWT), days to maturity (DM), pod weight (PDWT), pod width (PDWD), and pod length (PLN), was also used for correlation and cluster analysis **(Supplementary Table S6 and S7).**

### 2.2. Phenotypic evaluation for blanchability

For phenotypic assessment of groundnut blanchability, first, the regular groundnut grading was done where the variability in seed size observed from 7-8 mm diameter depending on the genotype. The primary objective of grading was to reduce the seed size and maturity variability. Subsequently, the pre-blanching weight of every genotype was measured. Then the phenotyping of groundnut germplasm was done, for obtaining the blanchability phenotypic data, seeds were subjected to hot air oven. Then the heated seeds were cooled to room temperature, for more than 8 hrs. The samples were treated on a blancher for sixty seconds per sample. According to the guidelines, seed sample weight of approximately 200 gm and 50 gm was pre-heated at 110°C for 30-35 min, the pre-heating results in reduction of moisture content to the range of 3.75-4.0 %. The samples were then allowed to cool down at room temperature. For medium size and extra-large seeds, the blanching time was determined to be 180±25 sec and 240±25 sec, respectively, under air pressure at 121±0.5 kPa (17.6 ± 0.1 psi) **(Figure 1)** [11]. In early generations, seed-set was low, thus a phenotyping technique that is based on small sample size is more desirable since it also offers a chance for high blanchability phenotypes in pedigree or in single-seed-descent programs [14]. Hence, in this study, we have considered two different sample sizes, 50 gm and 200 gm, with a blanching duration of 60 sec and 120 sec, respectively, at 30.00 rpm. The weight of the blanched seeds as well as blanched splits should be then observed and recorded **(Figure 2A)**, followed by calculating the blanching percentage, using the formula below:

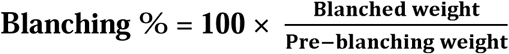

**Figure 1:**
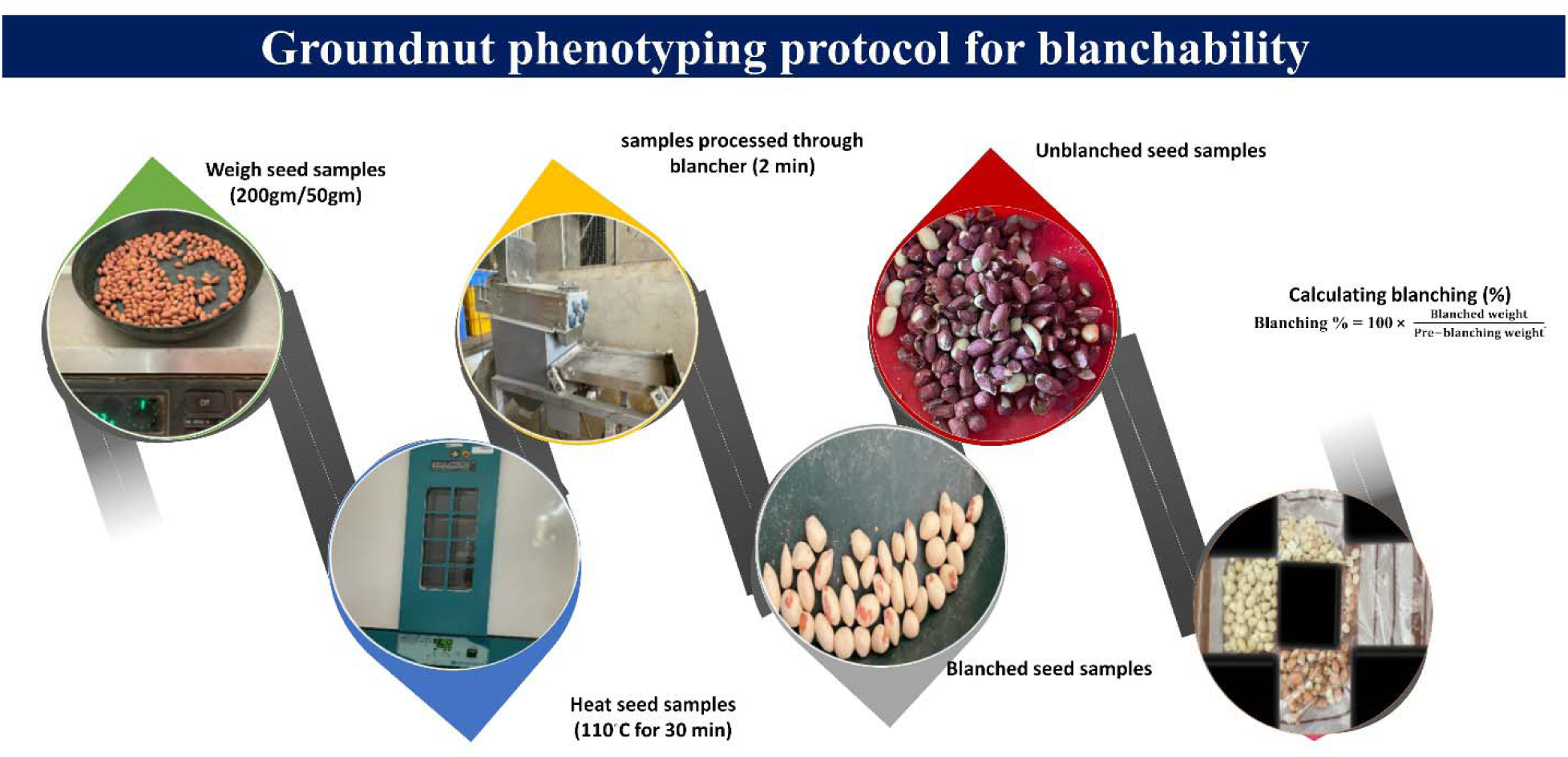
Protocol for phenotyping blanchability in groundnut

**Figure 2:**
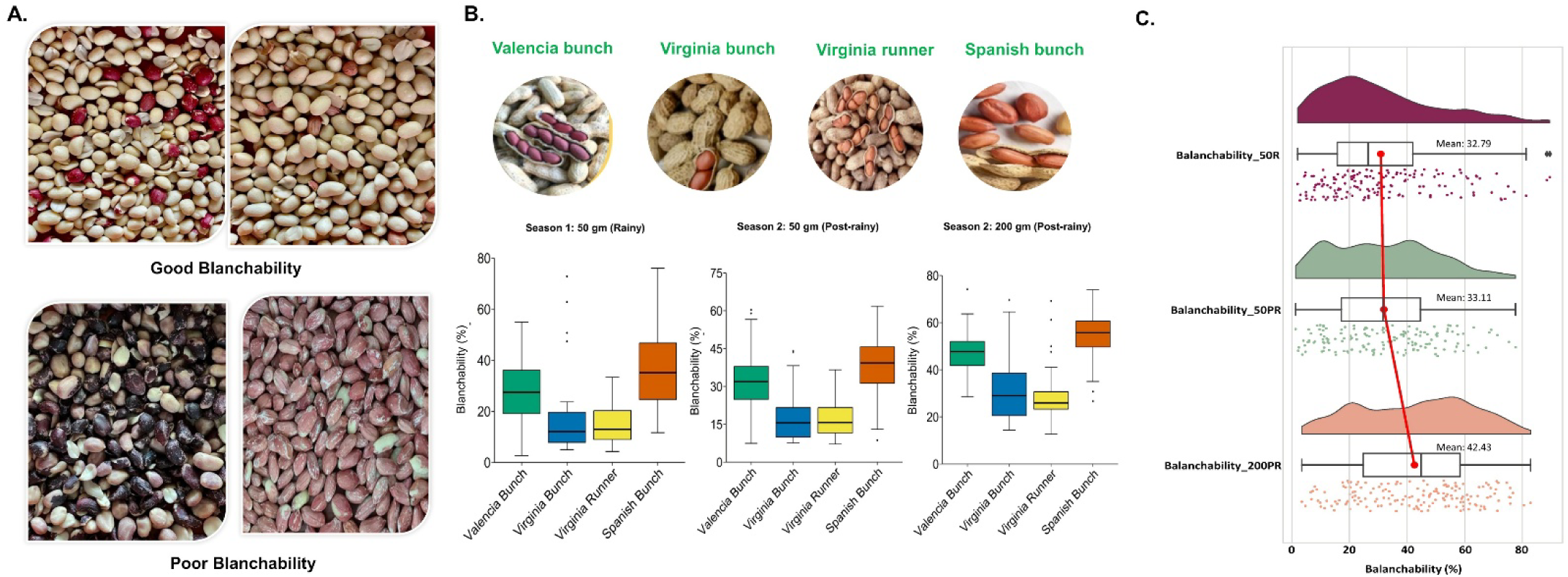
Different phenotypic analysis of blanchability across groundnut minicore: **A.** Phenotypic variability for blanchability in groundnut minicore collection, **B.** Diverse agronomic types in groundnut revealed substantial variation for blanchability in minicore collection across different seasons and sample sizes, **C.** The raincloud plot depicts the blanchability (%) values in season 1 (Blanchability_50R, pink color) and season 2 (Blanchability_50PR and 200PR, green and orange color; respectively), displaying considerable phenotypic variation with a continuous distribution of accession across both the seasons.

### 2.3. Phenotypic data interpretation using statistical analysis

The analysis of variance (ANOVA) and descriptive statistics was performed over the rainy (R) season of 2022 and post-rainy (PR) 2022-2023 using GenStat software version 15.0 (VSN International Ltd., Hemel Hempstead, UK) **(Supplementary Table S2, S3 and S4).** The phenotypic variation was observed in the minicore collection of groundnuts for blanchability trait was analyzed. In descriptive statistics, the Shapiro-wilk test was performed to test if the data follows normal distribution [24]. Analysis of differences between genotype replication through one-way ANOVA across different seasons was evaluated using the F-test test (0.05 significance level) **(Supplementary Table S2)**. Similarly, two-way ANOVA was used to study the genotype and environment interaction **(Supplementary Table S4).** The ‘GGplot’ package of RStudio (R Project for Statistical Computing) was used to analyze and plot the frequency curve on normal histograms and to plot the boxplot for showing distribution of phenotypic data across the agronomic types of groundnuts.

Pearson correlation analysis was conducted with customized RStudio script with ‘Hmisc’ package, and the correlation heatmap was generated with ‘pheatmap’ package in RStudio. Principal component analysis factor graph was employed to visualize the correlation among several traits, was calculated with ‘FactoMineR’ package in RStudio **(Supplementary Table S5 and S6, Figure 4)**. Minicore collection of 184 accessions was utilized for subsequent analyses. The frequency distribution plots were generated using the “ggplot2” package in the RStudio. The objective was to find a group of accessions to be utilized in breeding programs for good blanchability without compromising other agronomic qualities screened under various environments.

### 2.4. Principal Component analysis of phenotypic data

Principal component analysis was performed to examine phenotypic variability in terms of agronomic types and blanchability, and the contribution of these traits to the observed variation, for PCA analysis for each of these traits the phenotypic means were used. Further, based on this result of PCA, the clustering was done for phenotypic characters. The PCA was performed using FactomineR package of RStudio **(Figure 3 and Supplementary Figure S2).**

**Figure 3.**
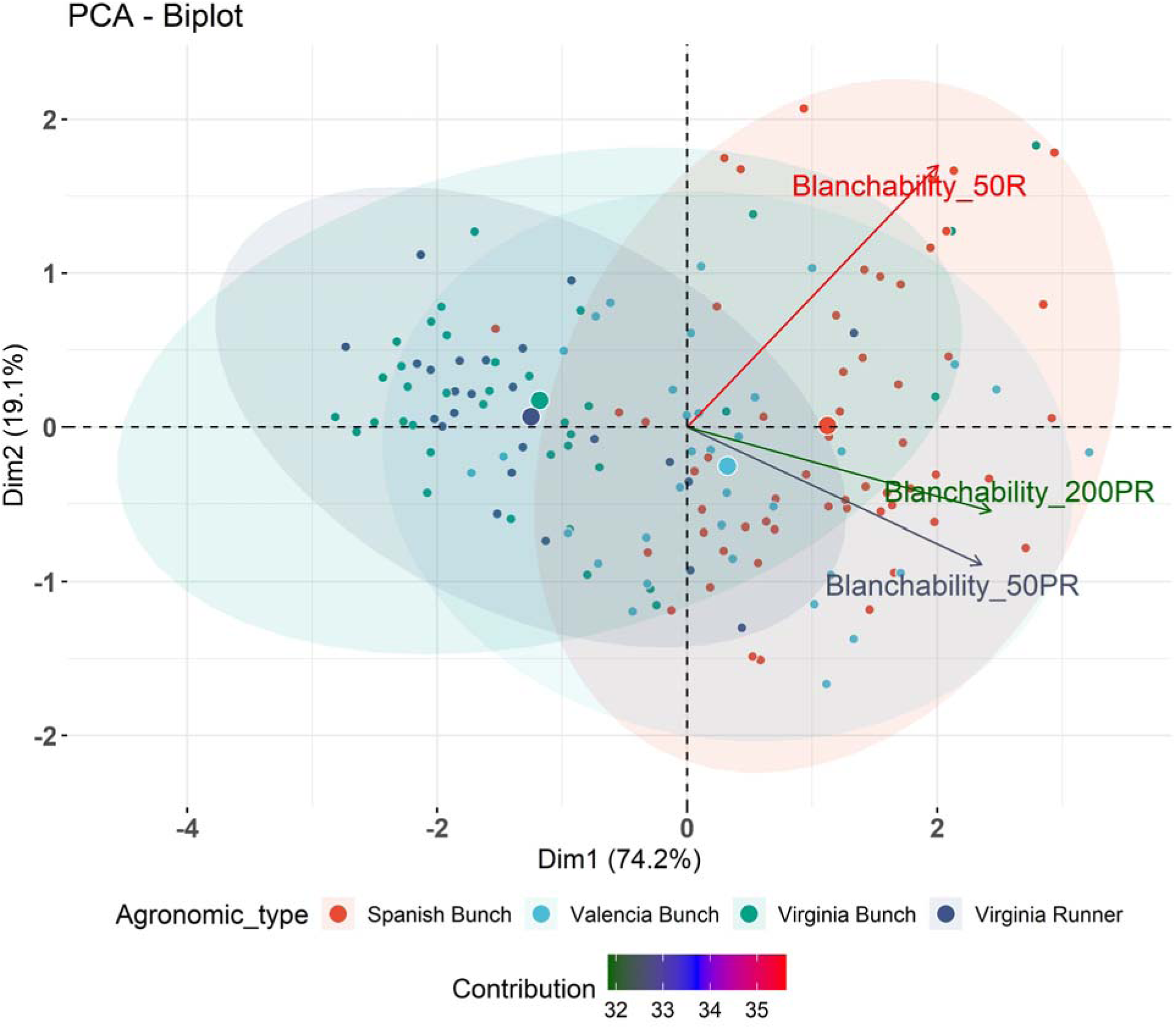
The PCA plot to depict the phenotypic variation. The distribution of the phenotypic dataset observed for blanchability across different seasons and sample size. The variation has been observed more for blanchability in 50 R. First two PCs collectively showing the 93.3% of phenotypic variation.

**Figure 4:**
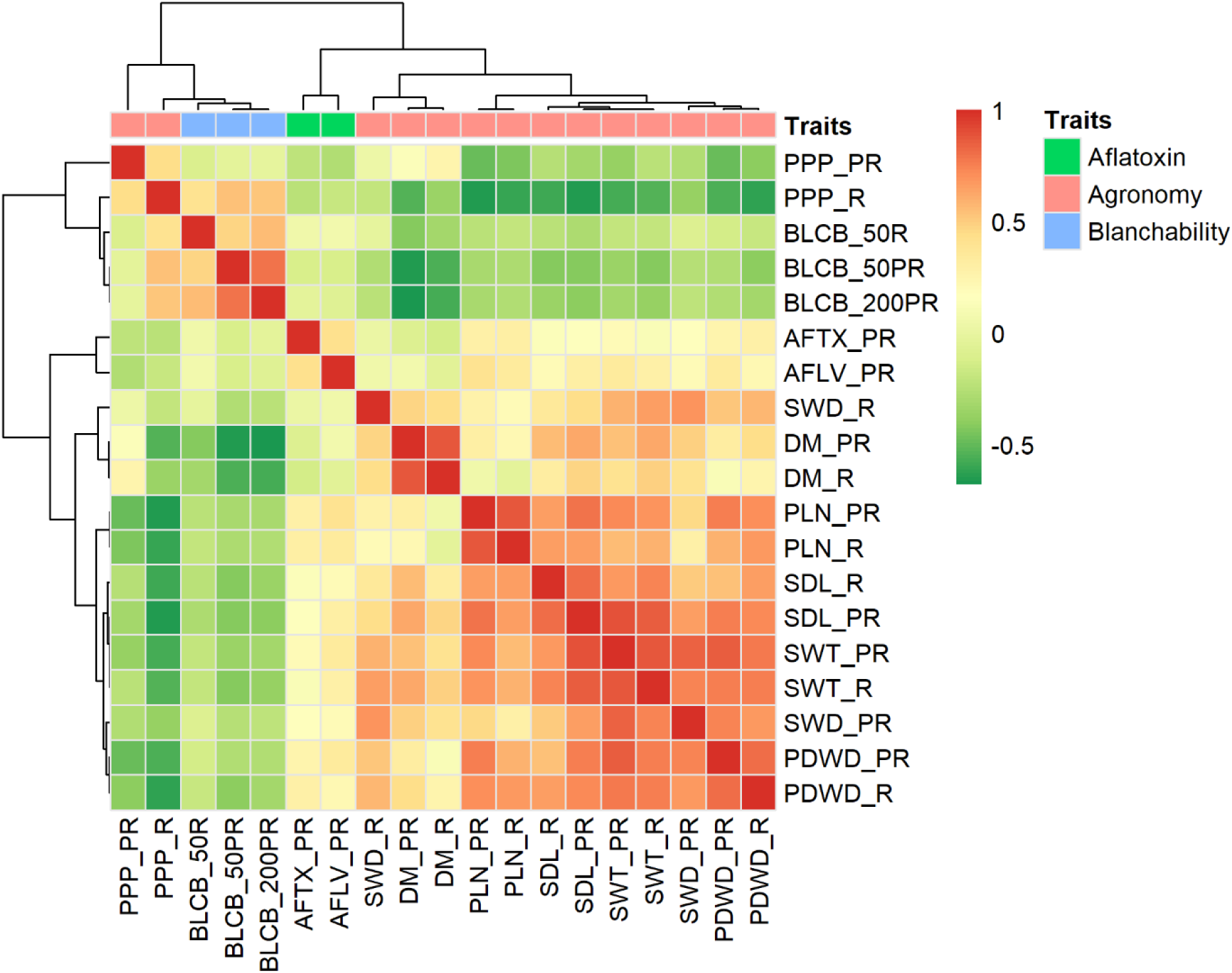
Pearson correlation analysis of blanchability, aflatoxin and agronomic traits were evaluated across diverse environments post-rainy (PR) and rainy (R). The four traits including pods per plant (PPP), blanchability (BLCB), aflatoxin (AFTX), *A. flavus* infection (AFLV), days to maturity (DM), pod length (PLN), seed length (SDL), seed weight (SWT), seed width (SWD), and pod width (PDWD) were evaluated in rainy and post-rainy environment. The color reflect the strength of correlation and further cluster has been made. PPP (R) positively correlated with blanchability and hence clustered together and other agronomic traits and aflatoxin make separate clusters. Scale ranges from -0.5 to 1, where the green color represents the least correlation while the red color shows strong correlation between the traits.

### 2.5. Identification of accessions as potential donors for blanchability

The accessions promising for blanchability and other agronomic qualities along with yield advantage, noticed in phenotypic screening at various environments, were chosen, based on favorable alleles. We analyzed the blanchability data produced in the current study and agronomic trait datasets to identify accessions that could serve as donors in the breeding programmes targeting high-yielding varieties with satisfactory blanchability. Two strategies were employed to select desirable accessions from the minicore collection, for good blanchability and agronomic performance across varying environments.

In this approach, the phenotypic data for blanchability and other agronomic traits, days to maturity (DM), seed length (SDL) seed weight (SWT), pod weight (PDWT), pods per plant (PPP), pod width (PDWD), pod length (PLN), were utilized in the identification of superior accessions. The accessions utilized here were based on trait correlations and hierarchical clustering **(Supplementary Figure S5)**. Three distinct clusters were formed and further then from each cluster to enhance the groundnut genetic potential superior accessions were identified to be used as potential donors in breeding programs. The basis of this selection was the presence of a favorable combination of at least two or more traits **[Supplementary Table S7 (a), (b), and (c)].**

### 2.6. 58 K ‘Axiom_Arachis’ SNP Array: A tool for high density genotyping in groundnut

A high-density assay of 58 K ‘Axiom_*Arachis*’ SNP Array [25] that encompasses genotypic information of 184 groundnut minicore accessions was utilized to find the SNPs-traits association. Genomic DNA was extracted from tender leaves of 25-30 days old seedlings of groundnut minicore collection accessions using Nucleospin Plant II kit (Macherey-Nagel, Düren, Germany). For genotyping, 20 ng/µl of each sample’s DNA was utilized with Affymetrix SNP array by the Affymetrix GeneTitan® system. Affymetrix GeneTitan® System was utilized as per protocol described by Affymetrix (Axiom® 2.0 Assay) and derived data for each accession was created and stored in CEL file format [25]. Genotyping was performed using ‘Axiom_*Arachis*’ SNP array that consists of 58,233 SNP markers that were obtained from DNA re-sequencing of 41 wild diploid ancestors and tetraploid accessions of groundnut [25, 26]. The 58 K SNPs correspond to an average of ∼2900 SNPs for each of 20 chromosomes.

Following SNP calling and data analysis conducted using Axiom™ Analysis Suite version 1.0 (Thermo Fisher Scientific, USA) to conduct quality control (QC) measures and choose samples that successfully cleared the QC test. SNPs were sorted into different classes followed by the “Axiom Best Practices Genotyping Workflow”. Polymorphic markers “Poly High Resolution”, and quality control passed samples were used in the study to assess the introgression lines with maximum recovery of the polymorphic SNPs. Monomorphic SNPs, low-intensity clusters, and SNPs having lower genotype call rates were removed while analysing the data.

TASSEL software was used to extract the polymorphic markers to further analyse the data. Of the 58,233 SNPs for 300 groundnut reference set were retrieved from Axiom™ analysis suit, the derived SNPs were further filtered for removing low quality and rare SNPs with ≤ 5% minor allele frequency (MAF) and >20% missing data using TASSEL (Trait Analysis by Association Evolution and Linkage) v5.2.93 software. After stringent filtration, high-quality SNPs across 167 accessions were subsequently employed for association mapping studies using GAPIT package in RStudio [27].

### 2.7. Population Structure analysis and genome-wide linkage disequilibrium

The population structure was estimated by PCA (principal component analysis), conducted using a genetic distance Roger’s matrix, to identify clusters of individual belonging to similar genetic background, which was calculated using selection tool package in RStudio **(Figure 5)**. The squared Pearson’s correlation coefficients (r^2^) for the pairwise analysis of SNPs was calculated to evaluate genome wise linkage disequilibrium (LD) decay [28]. LD decay in the groundnut minicore collection panel was calculated using PopLDdecay v.3.29. and selection tool package in RStudio **(Supplementary Figure S4 (B)).**

**Figure 5:**
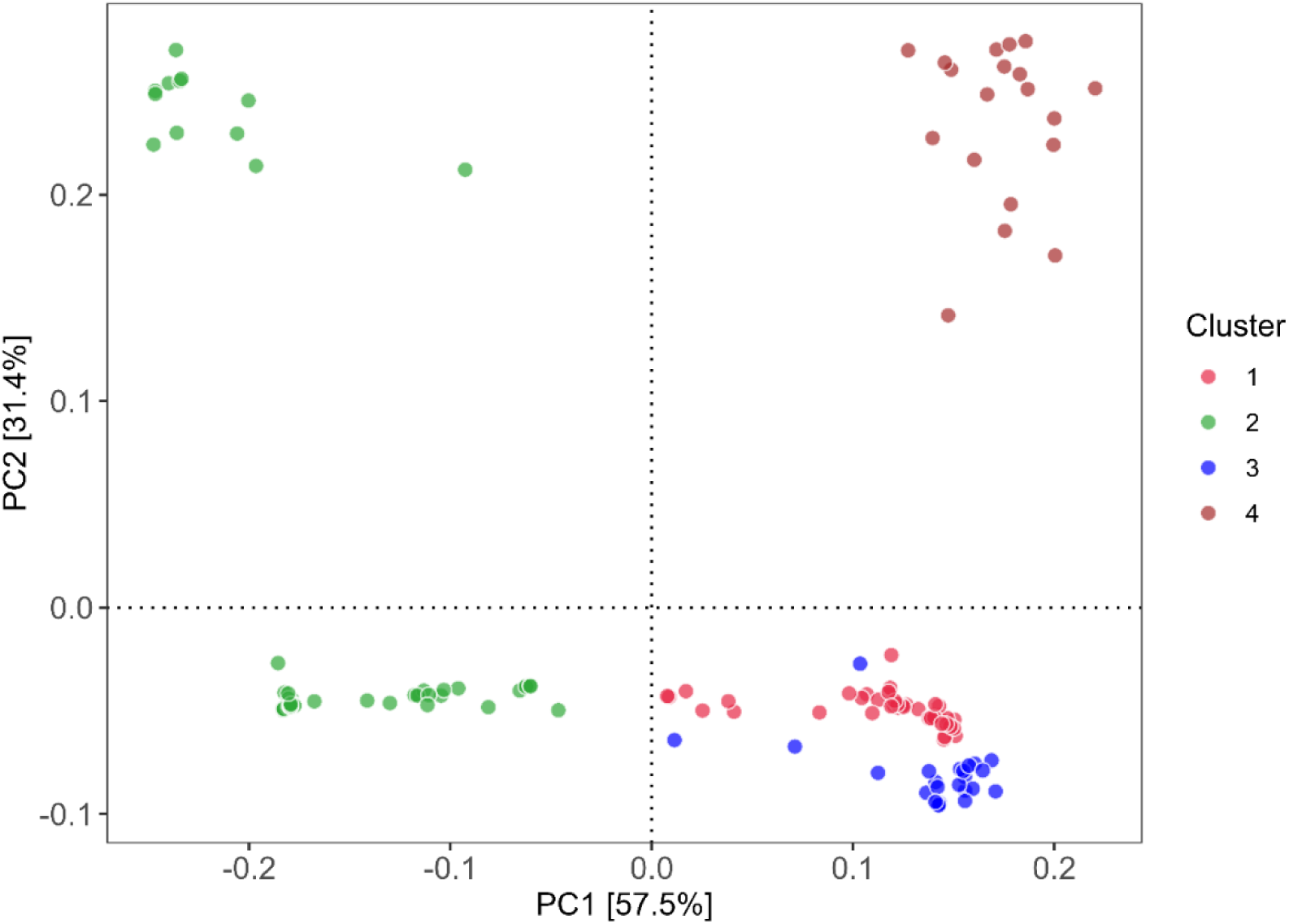
PCA depicting phenotypic variability of 88.9% for groundnut minicore collection, and identified 4 different clusters (pink, green, blue, brown) based on groundnut sub-species, botanical variety and agronomic type. Cluster 1 (pink color): (46 genotypes): *fastigata*, vulgaris, spanish bunch; Cluster 2 (green color): (75 genotypes): hypogaea, hypogaea, virginia bunch, Cluster 3 (Blue): (27 genotypes): fastigata, fastigata, valencia bunch, Cluster4 (Brown) (19 genotypes): fastigata, fastigata, vulgaris, spanish bunch, respectively.

### 2.8. Genome-wide association analysis for identifying SNPs-associated with blanchability

SNP-trait association was carried out utilizing advanced GAPIT models, namely, BLINK, FarmCPU, and SUPER that employed both population structure (Q) and kinship (K) matrices, utilizing GAPIT package of RStudio. Customize Quantile-Quantile (Q-Q) plots were created by graphing observed negative Log10 ‘p’ values against a computed expected negative Log10 ‘p’ value for all the SNPs available in CMplot package of RStudio [29]. A variation from ‘p’ values at the starting point reflects the population stratification present. Manhattan plots were employed to represent chromosome-wise SNPs acquired through the marker-trait association conducted throughout the genome. Log10 of ‘p’ values for every SNP were graphed against twenty chromosomes for the blanchability trait with the corresponding season and sample size. The false associations in GWAS were adjusted by applying “Bonferroni Correction” (95% confidence interval).

The threshold of association to establish significant SNP-trait association was calculated by applying Bonferroni correction for p-value of 4.9682 × 10-6, through negative log transformation of α/n (where, α refers to the overall significance level, n is the total number of SNPs employed for GWAS analysis). Phenotypic observations were taken on four sets of data: 200 gm (post-rainy), 50 gm (post-rainy), 50 gm (rainy), and 50 gm (average data of rainy and post-rainy). The percentage of phenotypic variance explained by all significant SNPs identified was reported from each model employed. The PVE% explained for each significant SNP was determined by the squared correlation between the SNP’s phenotype and genotype.

According to the distribution of SNPs, the threshold level of significance of associations between traits and SNPs was set at [–log 10 (*p*)< 10^-05^] for GWAS analysis by using SNP array that provided the best number of valid SNPs. SNP density plots, Q-Q plots, and Manhattan plots were created by the CMplot package of RStudio **(Figure 6, Supplementary Figure S1, S6 and Supplementary Figure S7) (Table 1).** In addition, as per the previously described methods by [30, 31], the impacts of the alleles behind the significant stable SNP markers were investigated. Genotypes were classified into independent groups based on their respective SNP alleles, and means were compared by Turkey’s HSD test **(Supplementary Figure S3).** In addition, the genes related to the STAs regions were found using the physical locations of significant SNPs in the reference genome sequence of Peanutbase (www.peanutbase.org). Gene function and effect of the corresponding SNPs were identified from the annotation file.

**Figure 6:**
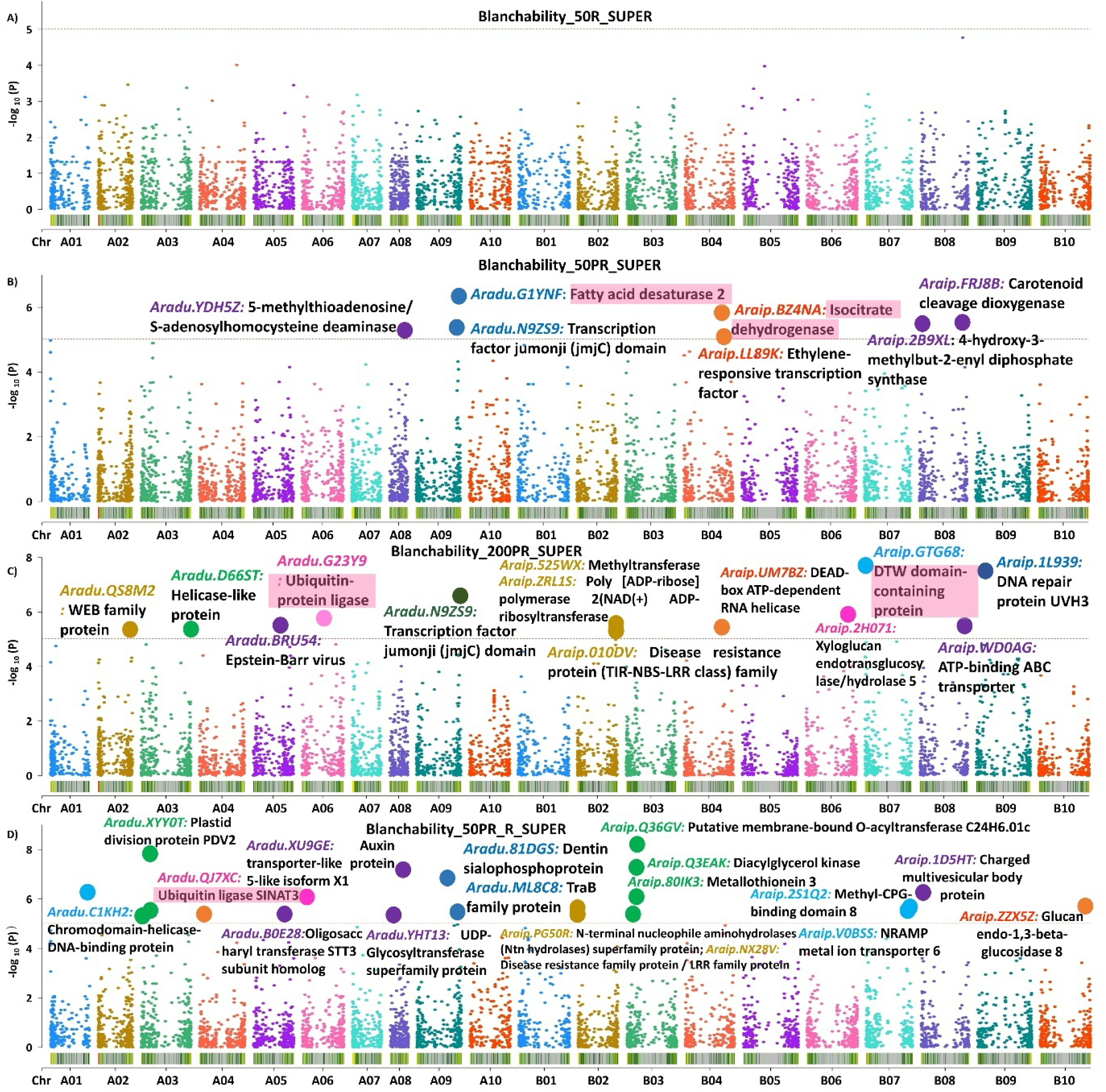
Manhattan plots representing the identified STAs associated with the blanchability on the basis SUPER model in GAPIT for. A: Blanchability 50R (50 gm sample size in rainy season), B: Blanchability 50PR (50 gm sample size in post-rainy season), C: Blanchability 200PR (200 gm sample size in post-rainy season) and D: Blanchability 50PR_R (50 gm sample size in post-rainy and rainy season), respectively. In these Manhattan plots, the highlighted (pink) region shows the gene associated with the validated polymorphic markers on chromosome A01, A06, B04 and B07; respectively. The other dots represent the significant STAs, and the genes found to be associated with blanchability.

**Table 1:**
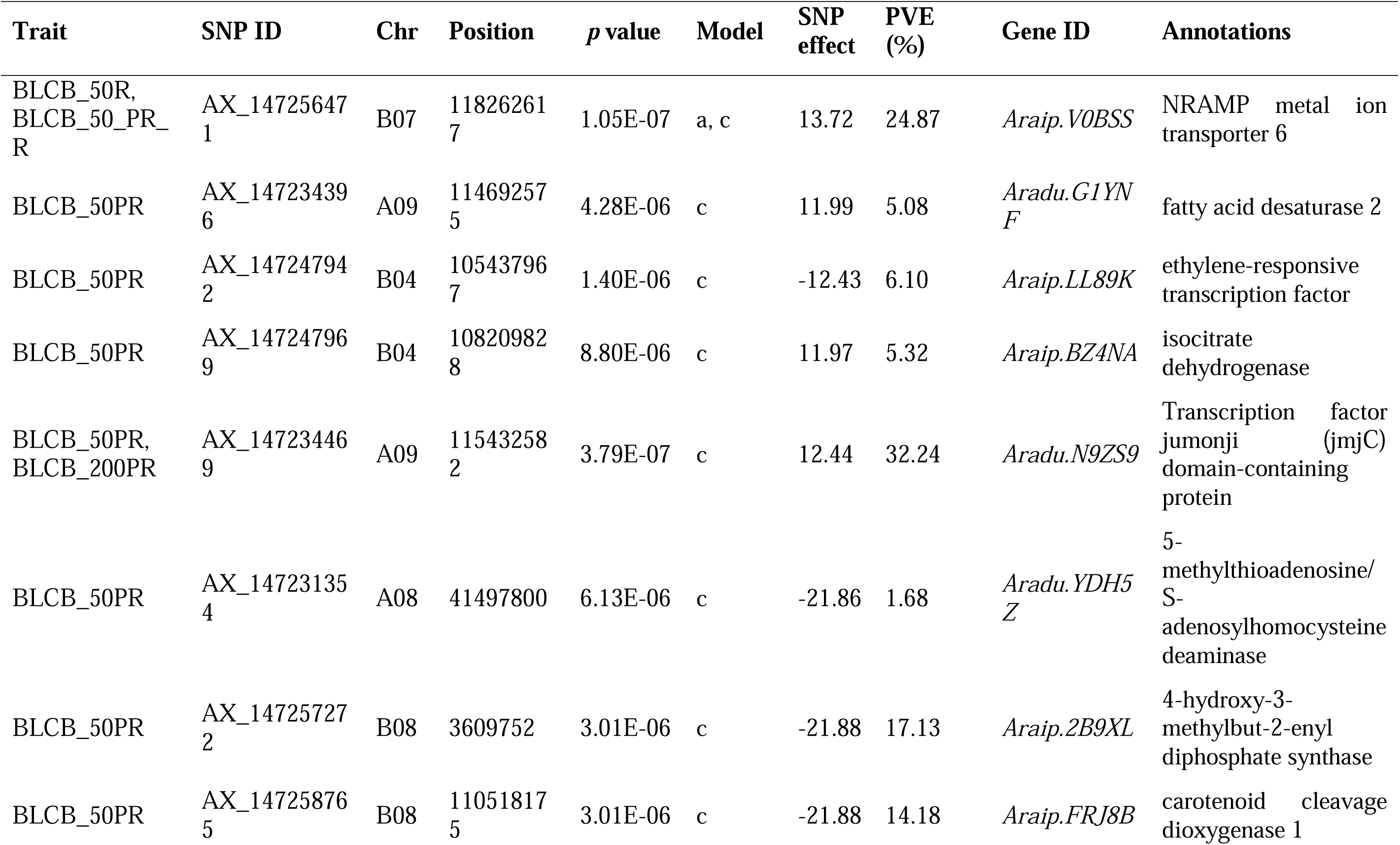
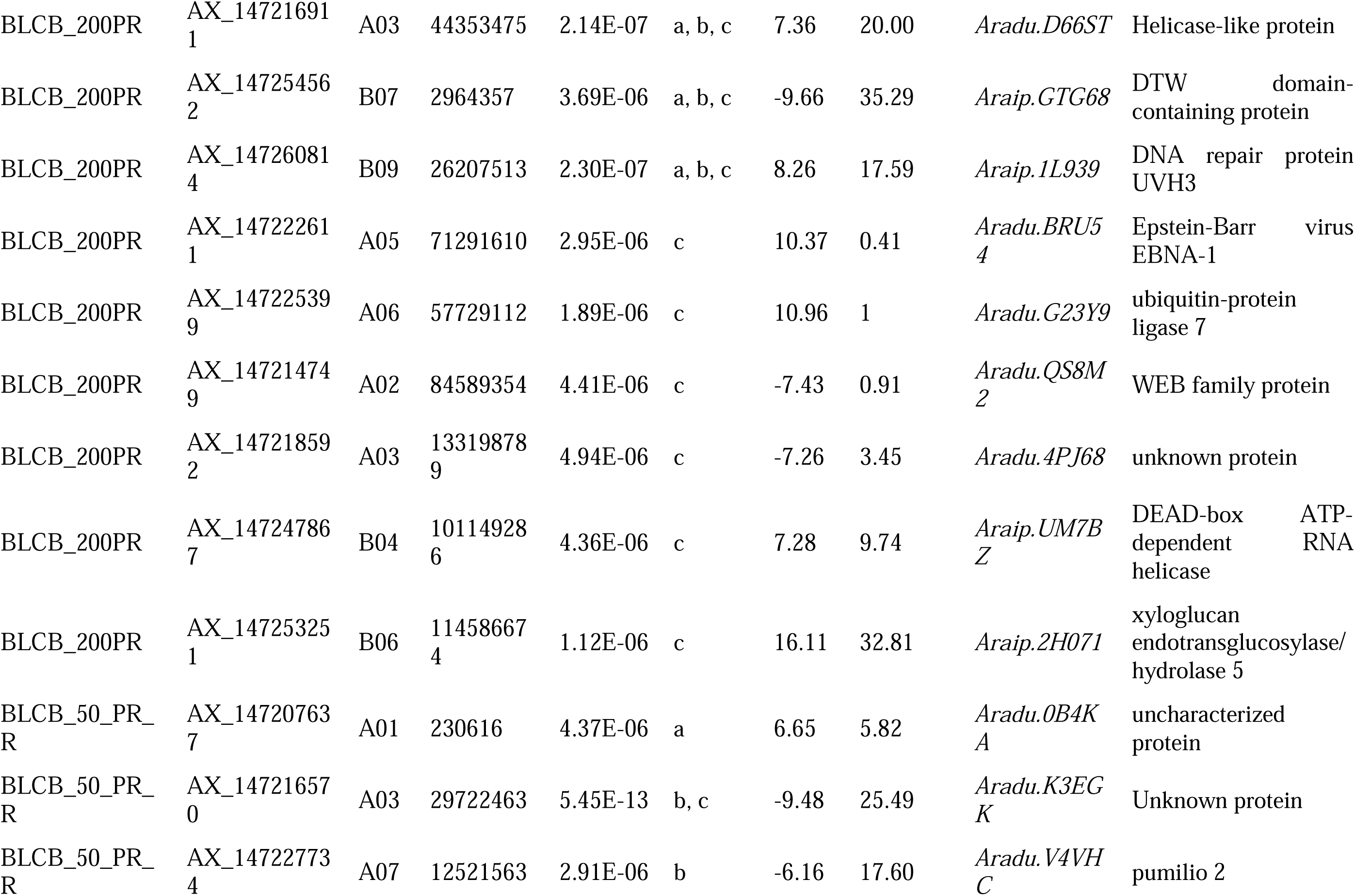
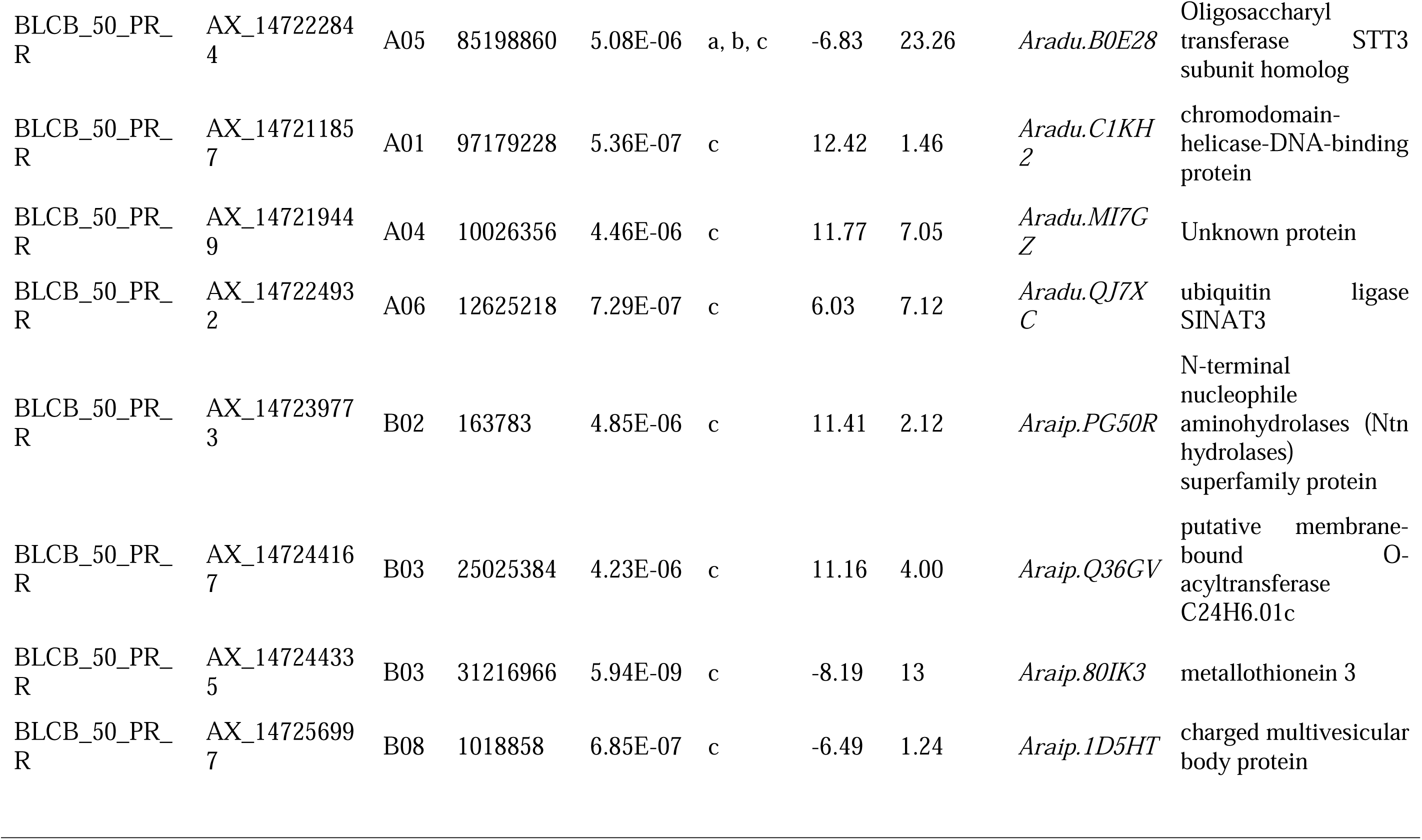
Marker trait association identified candidate genes and significant STAs for groundnut blanchability using 58 K “Axiom_*Arachis*” array

### 2.9. Candidate gene discovery associated with superior blanchability

Peanut base (https://Peanutbase.org/) was employed to detect the candidate genes using GBrowse (cultivated peanut) version 1. The SNP subsiding beginning and end position of the gene were investigated for the discovery of the candidate gene based on their biological function annotation towards the targeted trait **(Table 1).**

### 2.10. Development of Kompetitive allele specific polymerase chain reaction (KASP) -based markers

The present association mapping study identified 58 SNPs with a significant association, of which 10 highly significant markers were used for Kompetitive Allele-Specific PCR (KASP) assay development **(Supplementary Figure S8).** These SNPs were selected from genomic locations near candidate genes on three distinct chromosomes. The chosen SNPs were redesigned into KASP markers using 50-bp upstream and downstream regions to create user-friendly and economical markers [32]. Two allele-specific forward primers and a generic reverse primer were designed for every SNP marker (Intertek Pvt. Ltd.). The new KASP markers were validated across a panel of 25 genotypes, ranging from high blanchability and low blanchability genotypes as well as ICRISAT breeding lines **(Figure 7).**

**Figure 7:**
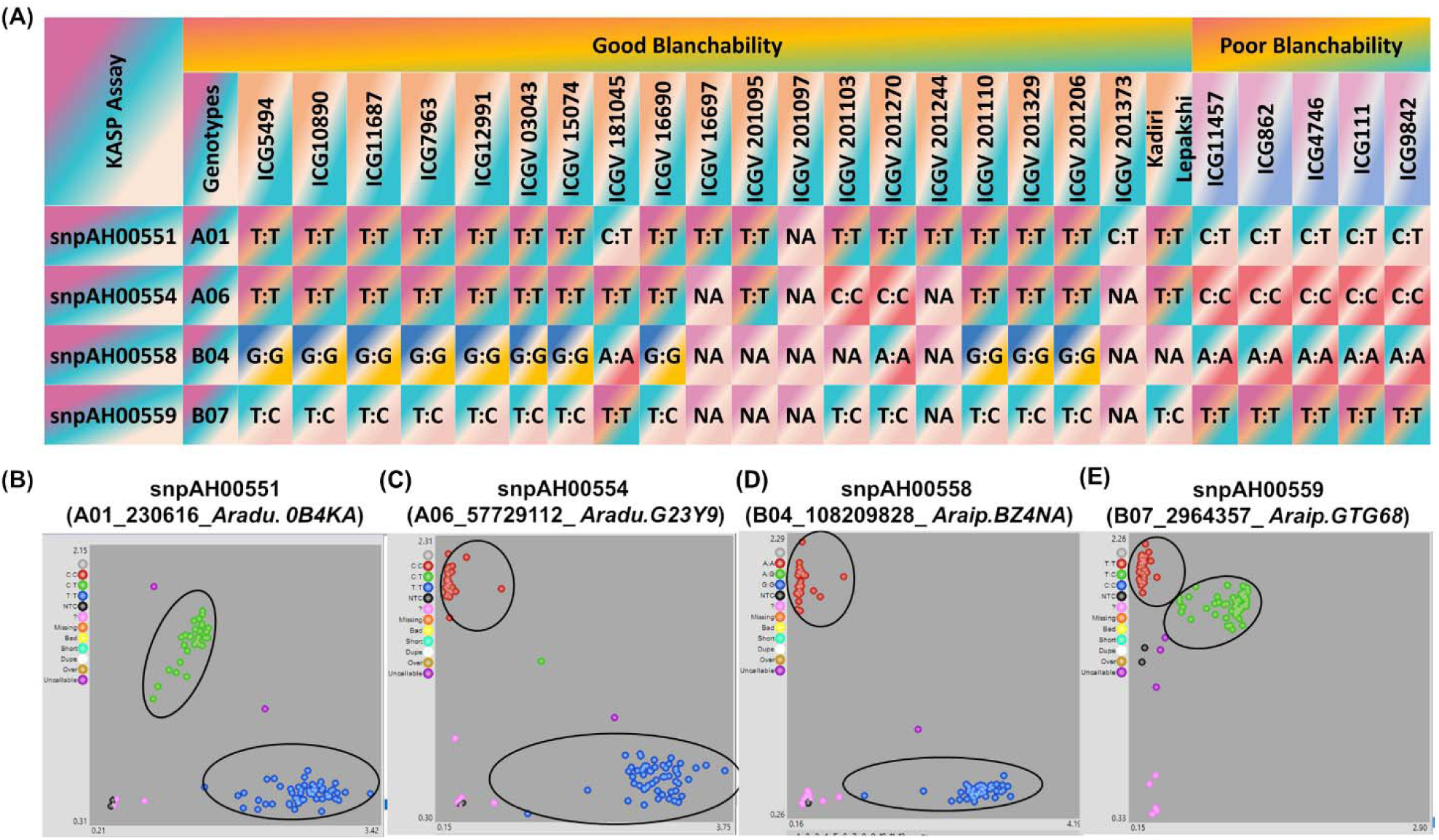
Development and validation of Kompetitive allele specific polymerase chain reaction (KASP) markers from potential candidate genes identified for blanchability. **A.** Validation panel includes 20 good blanchability genotypes and groundnut varieties (50-80%-blanchability), and five poor blanchability genotypes (1-10%-blanchability).Four single-nucleotide polymorphisms (SNPs) show clear homozygous clusters and heterozygous cluster for **B.** snpAH00551 (A01_230616) for uncharacterized protein (*Aradu.0B4KA*) gene, **C.** snpAH00554 (A06_57729112) for *ubiquitin-protein ligase* (*Aradu.G23Y9*) gene **D.** snpAH00558 (B04_108209828) for *isocitrate dehydrogenase* (*Araip.BZ4NA*) gene, **E.** snpAH00559 (B07_2964357) for *DTW domain-containing protein* (*Araip.GTG68*) gene.

## 3. Results

### 3.1. Phenotypic analysis of blanchability

Blanchability is one of the important economic traits and phenotypic variation is an essential component of GWAS analysis. The range and mean phenotypic distribution in 184 accessions from the groundnut minicore collection across different seasons and sample size are presented in **Figure 2C**. For season 2, traits exhibit a symmetric distribution, on the contrary, for season 1, with sample size 50gm, the distribution is slightly skewed to the right, indicating more values concentrated around 20 to 40 percent of blanchability, with fewer values beyond 60. **Supplementary Table S1** shows the wide variability range in the blanchability in the minicore collection. This includes blanchability_50R: 92.63 (mean 32.79), blanchability_50PR: 96.76 (mean 33.11) and blanchability_200R: 94.52 (mean 42.43). The trait and agronomic type wise phenotypic variation was depicted by box plot **(Figure 2B).** The agronomic type, spanish bunch and virginia runner, depicts the highest and lowest blanchability percentage; respectively. The blanchability trait was depicted across different seasons and sample sizes and it was observed that the variation has been profound in sample size 50 gm and rainy season.

Further, when the blanchability was correlated with the different agronomic types of groundnuts, Valencia Bunch, Virginia Bunch, Virginia Runner, and Spanish Bunch) across two seasons (Season 1: rainy and Season 2: post-rainy) and varying sample size (50 gm and 200 gm) **(Figure 2B).** The analysis determines that, in season 1 (rainy, 50 gm), spanish bunch has the highest blanchability, followed by valencia Bunch, while virginia runner shows the lowest. In season 2 (post-rainy, 50 gm), the general pattern remains, with spanish bunch still having the highest blanchability. However, with increased seed weight in season 2 (200 gm), blanchability increases overall, with the Spanish bunch maintaining its high blanchability and the other varieties showing a wider range of variation. The whiskers and outliers reflect variability within each variety **(Figure 2B).**

In the descriptive statistics analysis, it was observed that for rainy season (50 gm), values are generally lower and more around the mean, with a higher spread in the lower ranges, while post-rainy season (50 gm), the distribution is like the rainy season, though the mean is slightly higher. For post-rainy season (200 gm), values are significantly higher, with the mean around 42.57. Additionally, one-way and two-way ANOVA was performed using phenotypic data. In one-way ANOVA, it was observed that for the rainy season (50 gm), post-rainy season (50 gm), and post-rainy season (200 gm), the variation across genotypes was significant while the non-significant variation among replication suggests the accuracy of phenotypic data. In two-way ANOVA, the genotype and environment possess significant variation suggesting the effect of environment, while the genotype and sample size interaction is non-significant, that implies all genotypes respond similarly to sample size variations **(Supplementary Table S4).**

The PCA of the blanchability indicates that the PCs explained 93.3% of the total phenotypic variance **(Figure 3)**. The basis of clustering for accessions was not made on their agronomic type but had found to be varying degrees of relatedness, with Spanish bunch and Valencia bunch showing distinct separation, while Virginia bunch and Virginia runner overlap, indicating similarities. Blanchability_50PR and 200PR, were inversely related to the blanchability_50R. In addition, the PCA analysis of the blanchability and the agronomic traits showed that the first two PCs explained 62.3% of the total phenotypic variance. The analysis has found to highlight an interesting trend for the blanchability and the agronomic traits. AFTX_PR negatively related with PPP_PR, as did BLCB (200PR, 50PR, 50R) with DM (R and PR), and SWT have been found to highly significantly negatively correlated with BLCB (200PR, 50PR). In contrast, AFTX and AFLV are positively correlated with BLCB (200PR, 50PR, 50R) as did all other agronomic trait with BLCB (200PR, 50PR, 50R), however with AFTX and AFLV the relatedness is not significant **(Supplementary Figure S2)**.

These insights would find to be useful for targeting superior accessions in breeding programs. Pearson correlation analysis of blanchability, aflatoxin and agronomic traits were evaluated across diverse environments post-rainy (PR) and rainy (R). The four traits that include pods per plant (PPP), blanchability (BLCB), aflatoxin (AFTX), *A. flavus* infection (AFLV), days to maturity (DM), pod length (PLN), seed length (SDL), seed weight (SWT), seed width (SWD), and pod width (PDWD) were evaluated in rainy and post-rainy environment. The color reflect the strength of correlation and further cluster has been made. PPP (R) positively correlated with blanchability and hence clustered together and other agronomic traits and aflatoxin make separate clusters **(Supplementary Table S5 and S6) (Figure 4)**.

### 3.2. Potential donors for blanchability

Promising groundnut accessions with high productivity and blanchability can serve as donors for the breeding programs to be carried out to develop better varieties for fulfilling global demand. Employing information on blanchability and agronomic traits, for selecting better donors with favorable values of desirable traits. **Figure 4** indicates the correlation between blanchability and agronomic traits, where negative correlation exists for certain traits (days to maturity, seed weight). On the contrary, PPP (R) positively correlated with blanchability and hence clustered together and other agronomic traits and aflatoxin make separate clusters.

In identifying the possible donors in this strategy, we considered the correlations among the blanchability and agronomic characteristics, from various environments. All the characteristics describe nearly 62.3% of total variation. The PCA factor plot revealed that PC1 and PC2 explained 46.3% and 16% of the variation in measured characteristics, respectively **(Supplementary Figure S2).** Here, seed width (SWD_PR, SWD_R) and pod width (PDWD_PR, PDWD_R) were closely related to pods per plant (PPP_R) and pod length (PLN_PR, PLN_R) and distantly related to days to maturity (DM_R, DM_PR), similarly, blanchability (BLCB_50R, BLCB_50PR, BLCB_200PR) and seed weight (SWT_PR, SWT_R) are negatively correlated.

Hierarchical cluster analysis has made the classification of the traits into three distinct clusters: (1) days to maturity (DM_R, DM_PR); (2) seed width (SWD_PR, SWD_R), pod width (PDWD_PR, PDWD_R), pods per plant (PPP_R), seed length (SDL_PR, SDL_R); and (3) blanchability (BLCB_50R, BLCB_50PR, BLCB_200PR), seed weight (SWT_PR, SWT_R), pod length (PLN_PR, PLN_R), pods per plant (PPP_PR) **(Supplementary Figure S5).** The top accessions were selected based on phenotypic performance for the blanchability and agronomic traits. Accessions grouped in the same cluster were allowed to be compared among themselves, to identifying accessions for favorable combinations of two or more traits. For instance, cluster 1 had 21 accessions with higher days to maturity (**Supplementary Table 7 (a))**, twenty-seven accessions with cluster 2 had eight accessions that had higher SWD_PR, SWD_R, PDWD_PR, PDWD_R **(Supplementary Table 7 (b))**. In cluster 3, eight accessions had higher BLCB_50R, BLCB_50PR, BLCB_200PR, SWT_PR, SWT_R, PLN_PR, PLN_R, PPP_PR, and **[Supplementary Table 7 (c)].**

Selected intercrossing allows the development of improved accessions that have found to harbor beneficial alleles for blanchability and agronomical traits in groundnut. For example, ICG297 (cluster 1) could be crossed with ICG15419, ICG11219, ICG332, ICG6022, ICG9809, ICG 4729 and ICG11687 (cluster 3) to breed long-seeded groundnut varieties with good blanchability and seed weight. ICG6766 (cluster 2) could be crossed with ICG332 (cluster 1) to breed more days to maturity, seed width, pod width sand pod length varieties. We also made the comparison between the accessions identified in different cluster to identify common accessions between cluster 2 and 3, eleven accessions (ICG15419, ICG6022, ICG 11219, ICG4538, ICG11862, ICG6766, ICG9777, ICG4746, ICG2381, ICG8760 and ICG297), and accessions (ICG332) and (ICG 14630) common between cluster 1 and 3, and cluster 1 and 2, respectively. One accession (ICG297) was identified to be common between all the three clusters, cluster 1, 2 and 3, that could be used in breeding programs as potential donors to enhance both, groundnut blanchability and agronomic traits.

### 3.3. Genotyping and SNP density

The 58 K ‘Axiom_*Arachis*’ SNP array was used for the genotyping of minicore collection. After stringent filtration a total of 5044 high-quality SNPs across 167 accessions out of 184 as for some genotypes data was missing, this data was subsequently utilized for GWAS using GAPIT package of RStudio. For SNP array, SNP density across chromosomes, segmented into 1Mb windows, certain chromosomes, such as Chr A02, Chr A03, Chr A04, and Chr B03, show the highest overall SNP counts, with totals of 323, 293, 296 and 291 SNPs, respectively. In contrast, chromosomes Chr A10 and Chr B08 have relatively lower SNP counts; 198 and 206; respectively. (**Supplementary Figure S1**)

### 3.4. Population structure analysis and genome-wide linkage disequilibrium

Heatmaps and dendrograms of the kinship matrix indicated that there was clear clustering among the genotypes. The basis of this clustering was observed to be based on polymorphic SNPs for the studied genotypes, in the SNP array (**Figure 5) [Supplementary Figure S4 (A) and (B)] (Supplementary Table S8).** For SNP array, the population structure based on groundnut sub-species, botanical variety and agronomic type also revealed four distinct clusters: cluster 1 (46 genotypes): *fastigata*, vulgaris, spanish bunch; cluster 2 (75 genotypes): hypogaea, hypogaea, virginia bunch; cluster 3 (27 genotypes): fastigata, fastigata, valencia bunch; cluster 4 (19 genotypes): fastigata, fastigata, vulgaris, spanish bunch, respectively **(Supplementary Table S8).** The graphical representation of the LD characteristics of the minicore collection is presented for SNP array data in **Supplementary Figure S4 (B).** In the case of SNP array data, the mean r^2^ value for the genome was 0.12 and the LD decay was observed to start at an r^2^ value of 0.95 and half-decay at 0.15. The curve of LD decay cut off at 1Mbp which is the genome-wide critical distance to observe linkage. Therefore, markers linked to the same trait within this distance were classified as linked or associated.

### 3.5. Identifying SNPs and its associated candidate genes for blanchability

GWAS analysis was conducted based on 58,233 high-density SNPs having less than 20% missing data. They were scattered among the twenty groundnut chromosomes. SNP genotyping data of 58,233 SNPs together with population structure information and kinship matrix were utilized for conducting genome-wide association analysis for blanchability in the 2022 rainy season and 2022-2023 post-rainy seasons. The threshold level of -log10 *p* value was fixed at 5.0 for SNP array analysis above which the SNPs are reported to be significantly associated **(Figure 6)**.

A total of 58 highly significant STAs for blanchability were identified. Of the 58 STAs, 21 were identified for 200PR with -log10 *p* values ranging from 1.61 × 10^-8^ to 6.28 × 10^-6^ which explained 0 to 35.28% of phenotypic variation (PVE), and 28 were identified from 50PR_R mean data set, with -log10 *p* values ranging from 5.45 × 10^-13^ to 9.36 × 10^-6^ which explained 0 to 39.03% of PVE, further from 50R, 4 STAs were identified with -log10 ‘*p*’ values ranging from 1.18 × 10^-8^ to 1.18 × 10^-6^, and PVE range from 0 to 45.93% while for 50PR, 7 STAs were identified with -log10 ‘*p*’ values ranging from 3.79 × 10^-7^ to 8.80 × 10^-6^ and PVE from 0 to 32.24%. Finally, the 9 markers that were identified for blanchability from the analysis with the 4 different data set, and these were distributed across chromosome A01 (1), A05 (2), A06 (1), A09 (2), B04 (2), and B07 (1). Among these markers, B07_2964357 (AX_147254562) was observed to possess the highest value for phenotypic variation of 35.28% with a -log10 *p* value of 3.69 × 10^-**6**^ **(Table 1) (Figure 6).**

Groundnut genome sequencing accounted for genome size of 1.2 Gb in *A. duranensis* (36,734 genes) and for *A. ipaensis* it was 1.5 Gb (41,840 genes) and demystified the role of several genes for blanchability. Physical locations of each SNP marker of the current study were compared with the groundnut genome sequence to identify the gene function that underlies the corresponding SNP using Peanut base (https://Peanutbase.org/). Based on the previously reported genes documented in the literature, candidate genes which were previously reported to control seed coat integrity and adhesion were subsequently found to be candidate genes controlling blanchability. A total number of 13 SNPs were identified to possess candidate genes that were associated with blanchability **(Table 1)** viz., *Oligosaccharyl transferase STT3 subunit homolog (Aradu.B0E28), NRAMP metal ion transporter 6 (Araip.V0BSS); Transcription factor jumonji (jmjC) domain-containing protein (Aradu.N9ZS9); 5-methylthioadenosine / S-adenosylhomocysteine deaminase (Aradu.YDH5Z), fatty acid desaturase 2 (Aradu.G1YNF), ethylene-responsive transcription factor 7 (Araip.LL89K), isocitrate dehydrogenase (Araip.BZ4NA); 4-hydroxy-3-methylbut-2-enyl diphosphate synthase (Araip.2B9XL); xyloglucan endotransglucosylase / hydrolase 5 (Araip.2H071); putative membrane-bound O-acyltransferase (Araip.Q36GV), ubiquitin ligase SINAT3 (Aradu.G23Y9); N-terminal nucleophile aminohydrolases (Ntn hydrolases) superfamily protein (Araip.PG50R); charged multivesicular body protein (Araip.1D5HT)* were found associated with SNPs identified.

### 3.6. Allelic effects of stable STAs and validation of blanchability SNPs

A total of 9 significant STAs were observed to be polymorphic for the blanchability identified with SNP array analysis **(Supplementary Figure S8)**. All the nine SNP markers A01_230616 (50PR_R), A05_71291610 (50PR_R,200PR), A05_85198860 (50PR_R), A06_57729112 (50PR_R,200PR), A09_114692575 (50PR_R,50PR), A09_115432582 (50PR_R,200PR), B04_105437967 (50PR_R,200PR,50PR), B04_108209828 (50PR_R,50PR), and B07_2964357 (50PR_R,200PR), showed stable STAs with blanchability traits were further allowed to be used for determining the individual allelic effects on the studied traits. The identified alleles of these nine SNP markers were observed to have substantial effects on blanchability **(Supplementary Figure S3)**.

### 3.7. KASP markers development and validation for blanchability

KASP markers epitomized low-cost and less tedious genotyping assays to conduct early generation selection among breeding lines to indirectly select a specific phenotype [33]. For the development and validation of the diagnostic markers for blanchability, out of nine SNPs, six SNPs were located on A05, A09 and B04 with two SNPs on each chromosome, and three SNPs with one each on A01, A06, B07. These SNPS were substantially targeted for KASP markers development, Primers were then successfully designed and developed for 9 SNPs; A01_230616 (TT/CC), A05_71291610 (CC/TT), A05_85198860 (TT/GG), A06_57729112 (TT/CC), A09_115432582 (GG/AA), B04_105437967 (CC/TT), A09_114692575 (GG/AA), B04_108209828 (GG/AA) and B07_2964357 (CC/TT) and validated on a validation panel with contrasting blanchability. The validation panel comprised of good and poor blanchability genotypes and breeding lines ranging from 6% to 80%. Of the nine KASP markers selected for validation, four KASPs (snpAH00551, snpAH00554, snpAH00558 and snpAH00559) showed polymorphism in the designed validation panel. Interestingly, these four KASP markers were polymorphic and were clearly differentiating between genotypes that possess good and poor blanchability. These KASP markers were localized on genomic region detected on chromosome A01, A06, B04 and B07 **(Figure 7)**. The validated highly polymorphic KASP markers for good and poor blanchability genotypes and can be used as potential diagnostic markers for very early-stage selection of blanchability breeding material at the varietal development process.

## 4. Discussion

Groundnut is a globally cultivated food legume, celebrated for its high protein and unsaturated oil content. Blanchability quality becomes a significant trait regarding the development and utility of groundnut breeding lines for edible consumption [34]. In the research conducted so far for the blanchability trait, runner type groundnut has been utilized much more than other types and several laboratory-scale blanchers were customized and designed to assist breeding programs for the identification the high blanchability genotypes [35–38] . In the present study, a minicore collection panel was analysed using 58 K ‘Axiom_*Arachis*’ array and multi-environment phenotyping for the identification of the genomic regions and candidate genes that were associated with blanchability. This approach aims to enhance the understanding of the genetic factors influencing this economically important trait in groundnut cultivation.

In addition, some accessions identified through cluster analysis that can serve as potential donors and would further facilitate for the development of accessions with improved traits. The accessions would be harboring beneficial alleles governing blanchability along with agronomical traits in groundnut. Accession ICG297 (cluster 1) could be crossed with ICG15419, ICG11219, ICG332, ICG6022, ICG9809, ICG 4729 and ICG11687 (cluster 3) to breed long-seeded groundnut varieties with high blanchability and seed weight. ICG6766 (cluster 2) could be crossed with ICG332 (cluster 1) to breed high days to maturity, seed width, pod width and pod length varieties. Accessions (ICG297) between all three clusters, cluster 1, 2 and 3, combine blanchability and agronomic traits, that could be effectively used in breeding programs as potential donors. Therefore, these assays could significantly enhance future molecular breeding initiatives, providing a targeted and efficient approach to selecting lines with desired traits.

The cultivated agronomic type of groundnut includes, ‘Spanish Bunch’, ‘Valencia Bunch’, ‘Virginia Runner’, and ‘Virginia Bunch’. These botanical types have been found to possess vary distinct phenotypic characters, including branching habits, seed, and pod size which is according to the suitability to the specific cultivable environments. Spanish Bunch has been observed to possess good blanchability, while the maximum of virginia runner were with poor blanchability. So as per the industrial demand to produce groundnut processed food such as peanut butter, cookies, chocolate industries can prefer spanish bunch agronomic type (good blanchability), while for beer nuts and other seed based confectionary products virginia runner can be considered (Poor blanchability).

Genomics assistedbreeding has been an emerging technology for improvisation of several significantly important traits such as disease resistant, improving nutritional content, abiotic stress resistance in groundnut [39–41]. GWAS have become the primary method for identifying STAs related to complex traits of interest. Utilizing advanced models like FarmCPU, BLINK and SUPER in GWAS significantly enhances the reliability and precision of identifying genetic associations in plant genomics [42–44]. Resequencing-based genotyping reveals extensive natural variations, allowing for the exploration of functional genes and elite alleles within natural populations through GWAS. This approach enables us to identify valuable genetic traits that can enhance breeding programs and improve crop performance [45, 46]. In this study, we identified a total of 58 significant STAs for the blanchability trait using 58 K ‘Axiom_*Arachis*’ array; through GWAS using a minicore collection panel. Among which four SNP markers were validated, that were distinguishing the good (50-80%) and poor blanchability lines (1-30%), snpAH00551 (A01), snpAH00554 (A06), snpAH00558 (B04) and snpAH00559 (B07).

These results are supported by the previous study reporting the same chromosome to be associated with blanchability, however the genomic region varies [16]. The marker found in previous studies on A06 were located at *Arahy.06_108,665,514, Arahy.06 _108,812,907* which is located at a distant from the polymorphic marker, *Aradu.06_57,729,112*; we have found and validated in our study. These findings suggest that chromosome 6 (A06) comprises of an important region to be considered for blanchability. Additionally, the markers from chromosome A06 have also been validated by the Peanut Company of Australia Pty Ltd, A Bega Company, Australia.

Groundnut blanchability, is a trait influenced by genetic and environmental factors. Studies have associated blanchability with specific genomic regions or quantitative trait loci (QTLs). Previous research suggests that QTLs linked to groundnut blanchability are often identified on chromosome A06 and B01, these associations are determined through performing QTL-seq with two bulks of breeding lines from three populations. In addition, the KASP markers were developed from the most important SNPs from the QTLs on A06 *(Arahy.06_108,665,514, Arahy.06 _108,812,907) and B01 (Arahy.11_15,264,657 Arahy.11_16,329,544, Arahy.11_18,994,278*) [16]. The linkage-drag that’s reported to be associated with two prominent *A. cardenasii* introgressions (A02 and A03) have been conferred for disease resistance and negatively affects blanchability. As a result, when breeding for disease resistance using wild germplasm, there is a risk of co-introduction of undesirable traits such as reduced blanchability due to their close genetic linkage [16]. However, if there is a requirement of skin for processing application, then this linkage-drag could be proven to be beneficial such as in case of beer nut and other seed based confectionary products. As it would enable to select disease resistant and blanching resistant varieties.

Blanchability was neglected until now, however efforts have been in progress to identify the SNP-trait associations (STAs) associated with the high blanchability in the panel for diversity of characteristics such as blanchability of groundnut with an aim to develop parental varieties of groundnuts with enhanced blanchability in order to serve as donors of breeding programs groundnut improvement. The SNP allele analysis of the diverse minicore collection accessions for the identified significant STAs indicated that accessions carrying all favorable blanchability alleles tend to exhibit good blanchability. The validation panel for blanchability includes extreme genotypes representing good blanchability and poor blanchability genotypes, along with breeding lines. Those STAs which distinguish clearly good and poor blanchability lines hold promise for developing molecular markers, and accessions with all favorable good blanchability alleles can serve as donors in MABC programs aimed at achieving good blanchability lines. Among these, four markers snpAH00551 (A01), snpAH00554 (A06), snpAH00558 (B04) and snpAH00559 (B07) were successfully validated across minicore collection and groundnut breeding lines.

Blanchability in groundnuts is more commonly linked to factors such as, water absorption and seed coat integrity and adhesion, moisture content, and cell wall structure and biochemical components like pectin and lignin. Moreover, there are studies identifying other genes related to seed coat properties and adhesion, which can influence blanchability. The cell wall composition, pectin, cellulose, lignin, and hemicellulose are key components, while adhesion molecules include proteins involved in cell wall modification or strengthening. Seed coat adhesion and seed coat integrity are key factors influencing traits like blanchability, how tightly the seed coat adheres to the cotyledons, which are influenced by both mechanical properties and biochemical interactions.

Most of the genes found have expressions that are distant from the pathways typically involved in the synthesis and remodeling of cell wall components, which are more directly responsible for seed coat properties. The SNPs, snpAH00551 (A01), snpAH00554 (A06), snpAH00558 (B04) and snpAH00559 (B07) that were found to be polymorphic in the validation analysis are found to be associated with genes *ubiquitin protein-ligase (Aradu.G23Y9)* (A06) and *isocitrate dehydrogenase (Araip.BZ4NA)* (B04). These genes have found to have effect on cell wall dynamics influences the ease of seed coat removal, a key factor in groundnut blanchability. *Isocitrate dehydrogenase* produces α-ketoglutarate (α-KG), which regulates enzymes involved in cell wall loosening (e.g., *pectin methylesterases*, *expansins*). Changes in seedcoat integrity affect adhesion property, which is a determinant of blanchability [47].

Ubiquitin protein-ligase and ubiquitin ligase SINAT3 (a RING-type ubiquitin ligase) regulates protein degradation via the ubiquitin-proteasome system (UPS), affecting stress response, hormonal signalling, and cell wall modifications. SINAT3 is involved in abiotic stress response pathways, including oxidative stress and hormone signalling (e.g., auxin, ethylene), which regulate seed coat integrity and blanchability by influencing pectin breakdown and adhesion strength [48]. It likely interacts with transcription factors (e.g., NAC, MYB) involved in cell wall degradation, modulating lignin biosynthesis and pectin remodelling [49].

The interplay between oxidative metabolism (*Isocitrate dehydrogenase*) and protein turnover (*Ubiquitin protein-ligase* and *Ubiquitin ligase SINAT3*) determines seed coat adhesion strength, thereby influencing blanchability. *Isocitrate dehydrogenase* contributes to blanchability by regulating α-KG-dependent enzymes involved in seed coat loosening. *Ubiquitin protein-ligase* and *Ubiquitin ligase SINAT3* impacts blanchability by targeting proteins that control seed coat adhesion and degradation. This pathway can be further validated by transcriptomic and proteomic analysis of high- and low-blanchability groundnut genotypes to assess differential expression of *ubiquitin protein-ligase/ubiquitin ligase SINAT3*, *isocitrate dehydrogenase* and cell-wall-modifying enzymes.

If *isocitrate dehydrogenase* activity is high, lignin/pectin biosynthesis is enhanced, leading to a stronger seed coat and lower blanchability. If SINAT3 degrades key regulators of cell wall biosynthesis, this can lead to a weaker seed coat and higher blanchability. *Isocitrate dehydrogenase* affects blanchability via metabolic flux regulation, providing intermediates for seed coat biosynthesis. SINAT3 influences blanchability through the ubiquitin-proteasome system, modulating stress responses and cell wall composition. Their interaction is likely mediated by oxidative stress pathways, hormonal regulation, and protein degradation, ultimately impacting seed coat adhesion and blanchability in groundnuts.

Furthermore, we have determined genes that are associated to cell wall biosynthesis, modifications, remodeling and adhesion proteins such as *xyloglucan endotransglucosylase*, and *oligosaccharyl transferase STT3, putative membrane-bound O-acyltransferase* that influence seed coat’s biochemical properties. For instance, x*yloglucan endotransglucosylase (XETs),* participated in cell wall modification and maintained cell wall plasticity, helping plant in tolerating stress such as salinity stress [50].

*XETs* are enzymes that modify xyloglucans, major hemicellulosic components of the plant cell wall. They play a pivotal role in restructuring the cell wall by cleaving and rejoining xyloglucan chains, facilitating cell wall loosening and expansion. This activity is crucial for maintaining cell wall integrity and modulating adhesion between cells, directly impacting plant growth and development. Their function is vital during seed coat formation, where precise cell wall modifications are necessary for proper development [51].

The roles of *STT3* are a component of the *oligosaccharyltransferase (OST)* complex, which is involved in N-linked glycosylation, and glycoproteins in the cell wall or membrane could influence cell wall remodeling or stability and *stt3a* mutation disturbs the essential adaptation mechanisms [52]. *STT3* is a catalytic subunit of the *OST* complex, responsible for transferring oligosaccharides to nascent proteins during N-linked glycosylation. Proper glycosylation is essential for the stability and function of many cell wall proteins, thereby influencing cell wall integrity and cell-cell adhesion.

The structure of the seed coat is tightly regulated during seed maturation, and fatty acids, synthesized and modified by *Fatty acid desaturases (FADs) FADs*, are part of the cuticle (a waxy protective layer) that forms on the outer surface. In seed coats, the fatty acid profile affects the permeability and protective functions of the seed coat, thereby influencing seed development and viability [53]. *FADs* play a vital role for determining the lipid composition of the seed coat, affecting its physical properties and functions, including protection, permeability, and regulation of dormancy and germination.

All these enzymes are integral to the dynamic processes of cell wall construction, modification, and maintenance, directly affecting seed coat development, cell wall integrity, and adhesion in plants. By modifying interactions within the cell wall, these genes may influence how tightly the seed coat adheres to the cotyledons. Strong adhesion can make blanching more difficult, while effective modification might facilitate easier removal of the seed coat.

## 5. Conclusion

Blanchability is a trait of tremendous economic importance as there is a significant energy, time and cost requirement to remove the groundnut seedcoats from poor blanching lines varieties. Product processing ultimately becomes cumbersome leading to re-processing. This ultimately leads to considerable loss of kernels during the processing and sorting of the groundnut. The present study utilized 58 K ‘Axiom_*Arachis*’ array genotypic data from a diverse minicore collection panel phenotypic data collected over two seasons, to conduct a genome-wide association study. This phenotypic analysis reveals that the spanish bunch agronomic types are possessed to have good blanchability, additionally, the statistical analysis has revealed the effect of sample size and environment on blanchability trait. ICG297 genotype has been identified to be the superior potential donor for blanchability along with other agronomic traits. Further, the GWAS analysis has identified significant STAs in 58 K ‘Axiom_*Arachis*’ array genotypic data for blanchability. The analysis of these STAs revealed potential candidate genes such as *isocitrate dehydrogenase* and *ubiquitin ligase protein* associated with blanchability, highlighting the critical role of these genes in the dynamic processes of cell wall construction, modification, and maintenance and have a direct impact on the development of seed coats, the integrity of cell walls, and plant adhesion. Ultimately, four KASP markers snpAH00551 (A01), snpAH00554 (A06), snpAH00558 (B04) and snpAH00559 (B07) were validated from 58 K ‘Axiom_*Arachis*’ array exhibiting clear polymorphism between good and poor blanchability genotypes were identified, to be incorporated in breeding programs. Among these A06 chromosome, has been found common with previous study which suggests it could prove to be the future potential marker for blanchability. Furthermore, conducting haplotype analysis on these candidate genes could facilitate the identification of superior haplotypes for blanchability, which can be leveraged in haplotype-based breeding strategies.

## Data availability statement

The phenotypic data used in this work is provided in Supplementary Table S1. The sequencing data generated in this study is deposited in NCBI with bio-project ID PRJNA1002116.

## Institutional Review Board Statement

Not Applicable

## Informed Consent Statement

Not Applicable

## CRediT authorship Contribution Statement

**Manish K. Pandey:** Conceived the idea and supervised and finalized the manuscript. **Kuldeep Singh and Ramachandran Senthil:** Contributed seed material and in seed multiplication of the minicore collection. **Pasupuleti Janila:** Contributed the breeding lines for marker validation. **Priya Shah:** Phenotyped minicore collection, performed the analysis and wrote the manuscript. **Sunil S. Gangurde, Prashant Singam and Sean Mayes:** Reviewed, edited and improvised the manuscript. All authors have read and agreed to the published version of the manuscript.

## Acknowledgments

Priya Shah acknowledges Joint Council of Scientific and Industrial Research-University Grant Commission (CSIR-UGC), Government of India for the award of fellowship for a Ph.D. and ICRISAT for providing research facilities. The authors are thankful to Khaja D. Mohinuddin, for his assistance in improving manuscript figures.

