## Supplementary table S1-S8 and Supplementary Figures S1-S8 for "Identification of High Blanchability Donors, Candidate genes and Markers in Groundnut": Supplementary.Figures.docx

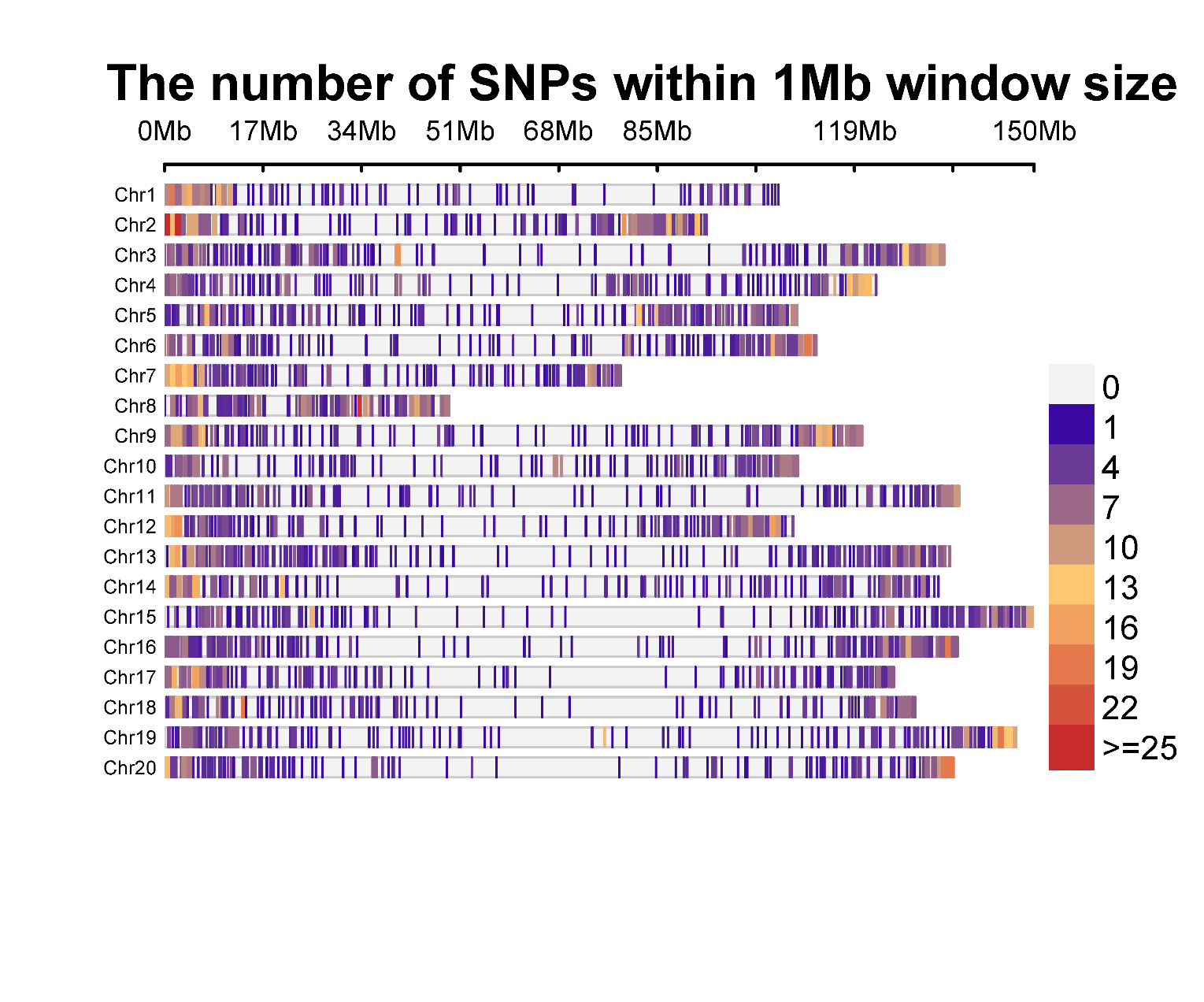

**Supplementary Figure S1: SNP density plot with 1Mb window size for 58 K ‘Axiom_*Arachis*’ array data.** Red-colored regions, indicating the highest SNP density, appear sparsely, particularly on Chr A02. There are also white gaps on chromosomes, signifying regions with low or no detected SNPs.

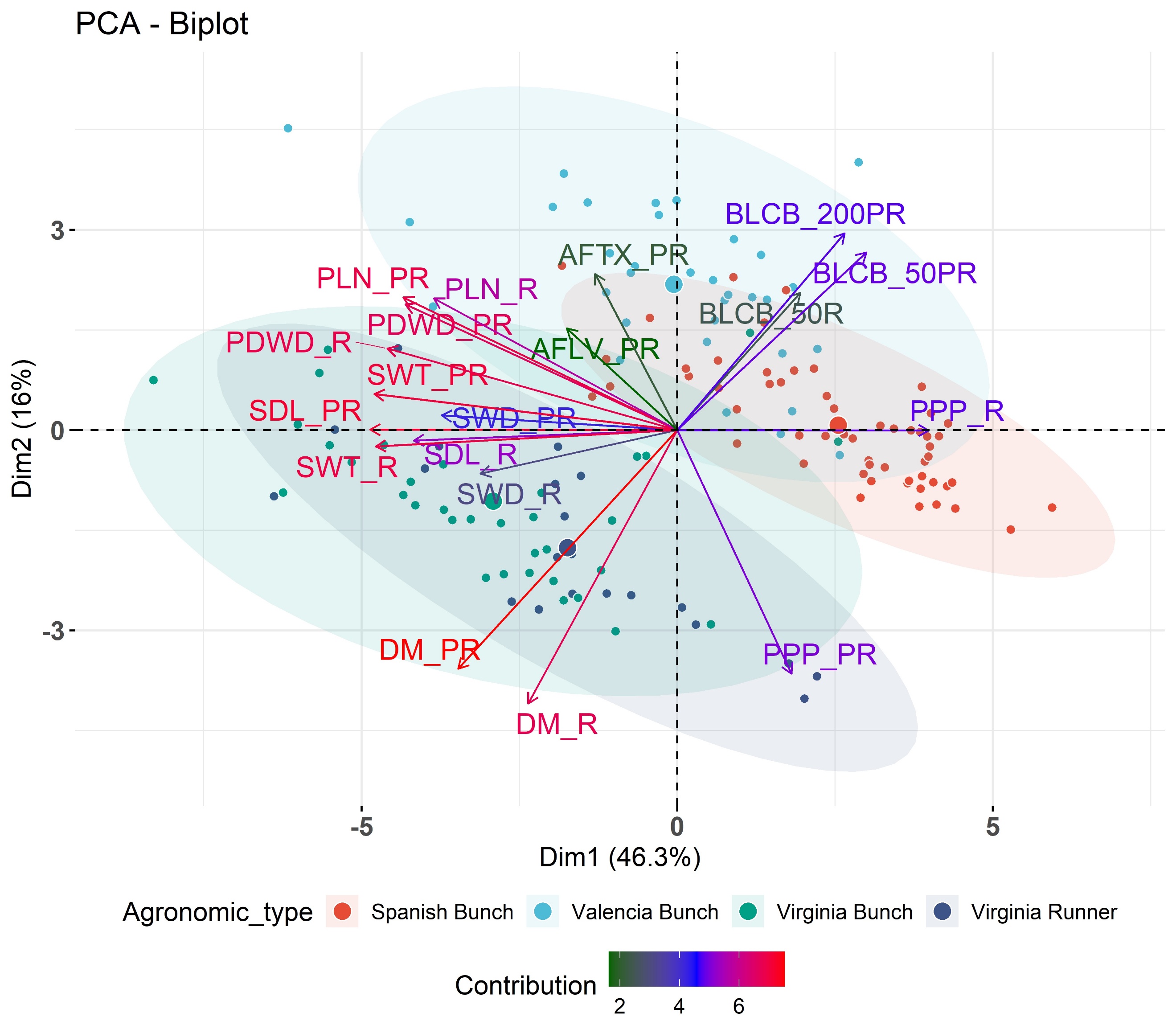
**Supplementary Figure S2. Principal component analysis for blanchability, aflatoxin and other agronomic traits.** Projection of the minicore collection on the first plane of principal component analysis using phenotypic data for blanchability, aflatoxin and other agronomic traits [DM (day to maturity), SWD (seed width), PPP (pods per plant), PDWD (pod width), SDL (seed length), SWT (seed weight), BLCB (Blanchability) (50 gm) (200 gm), PLN (plant length) (PR:post-rainy and R:rainy)] across different season and sample size. The first two components, PC1 and PC2, explain 62.3% of the variance between genotypes. Among the traits, BLCB and DM have been observed to be negatively associated.

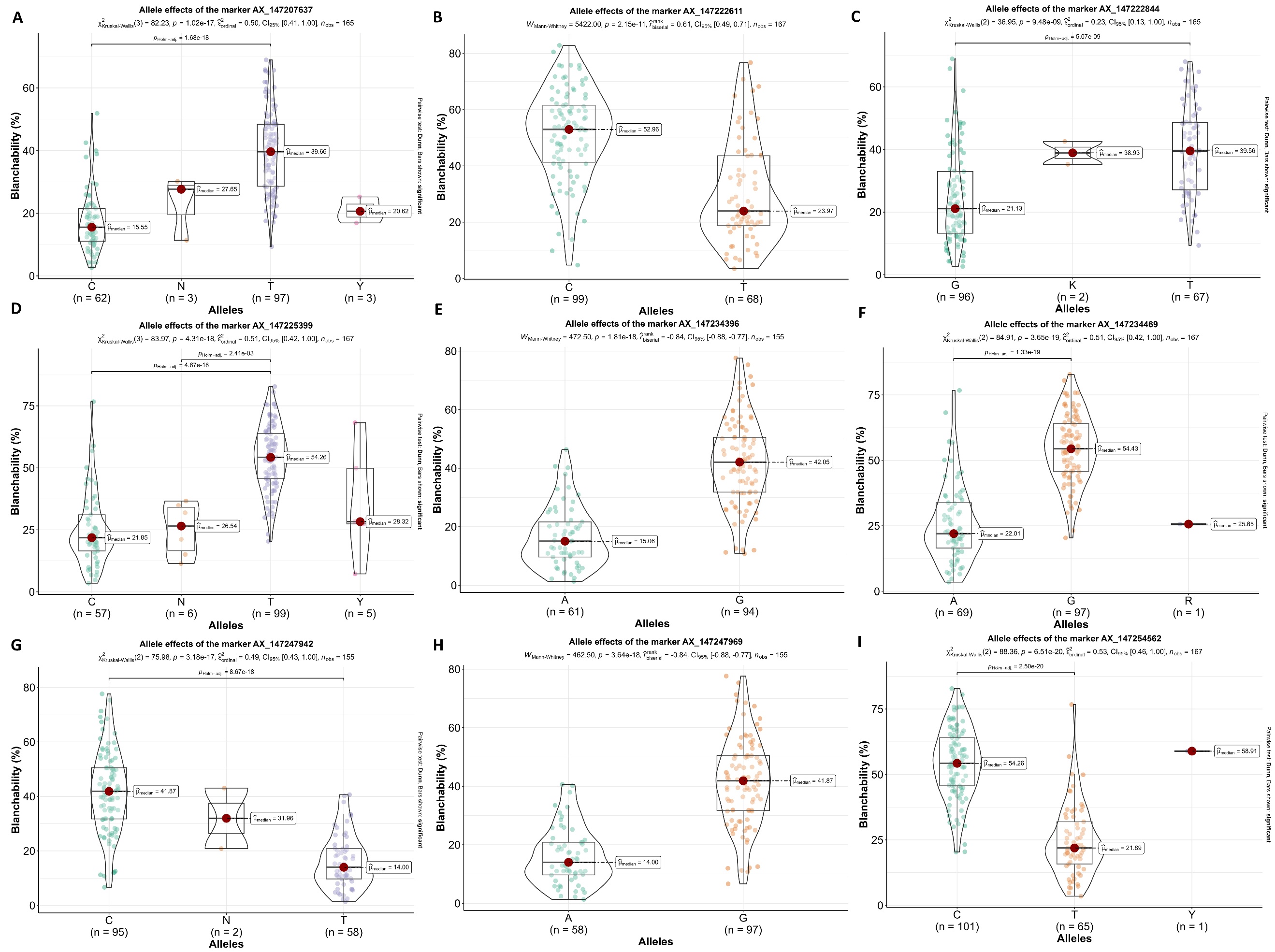

**Supplementary Figure S3: Allele-effect analysis for nine stable signiﬁcant SNPs** including **A.** AX147207637(A01), **B.** AX147222611(A05), **C.** AX147222844(A05), **D.** AX147225399(A06), **E.** AX147234396(A09), **F.** AX147234469(A09), **G.** AX147247942(B04), **H.** AX147247969(B04), **I.** AX147254562(B07). The plot depicts the number of the alleles for each of the nine signiﬁcant SNPs in minicore collection, and the contribution of these alleles to the phenotypic variation observed blanchability**.**

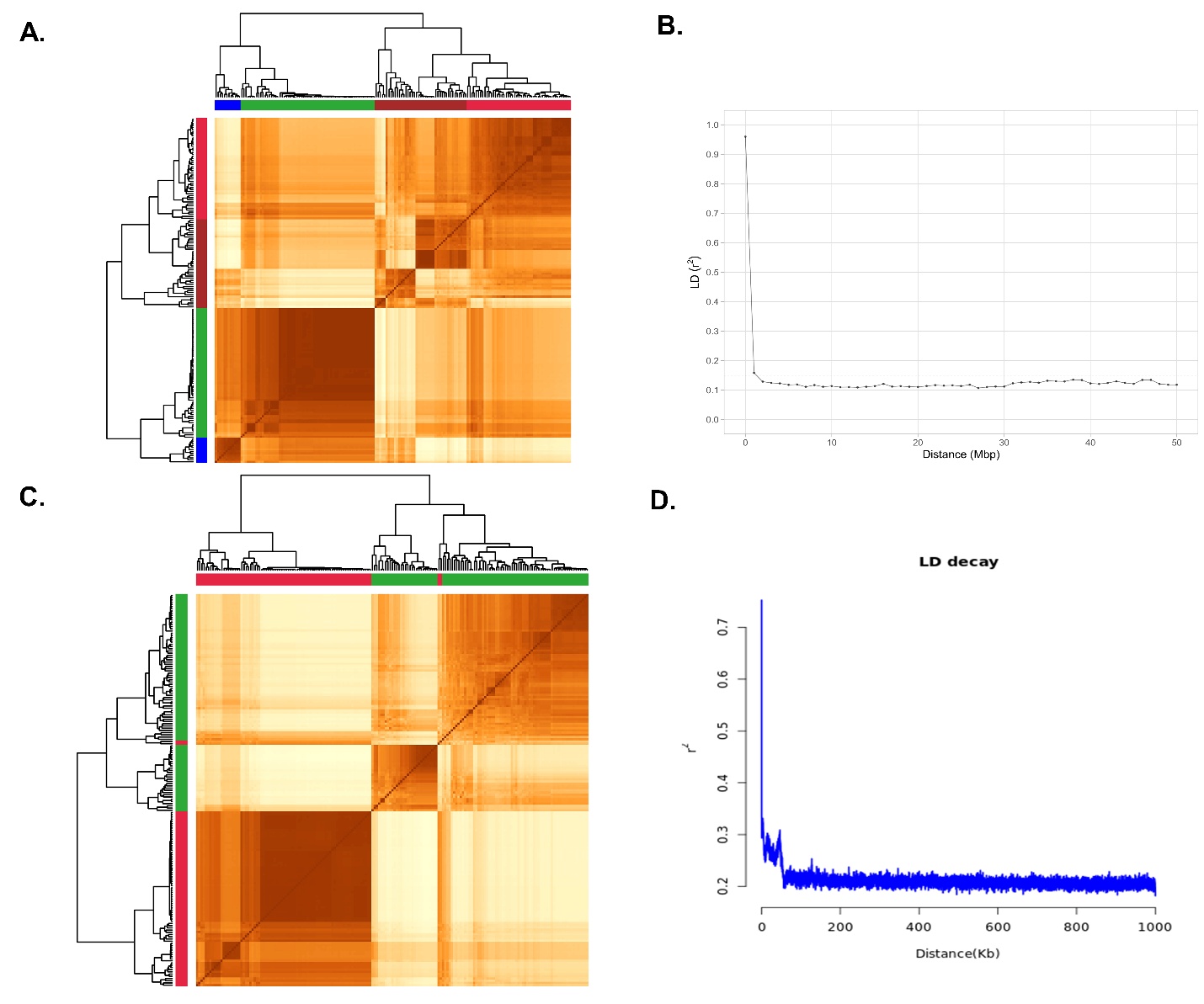

**Supplementary Figure S4: Population Structure analysis and genome-wide linkage disequilibrium, A.** Heat map of population structure, **B.** LD decay for groundnut minicore collection identified.

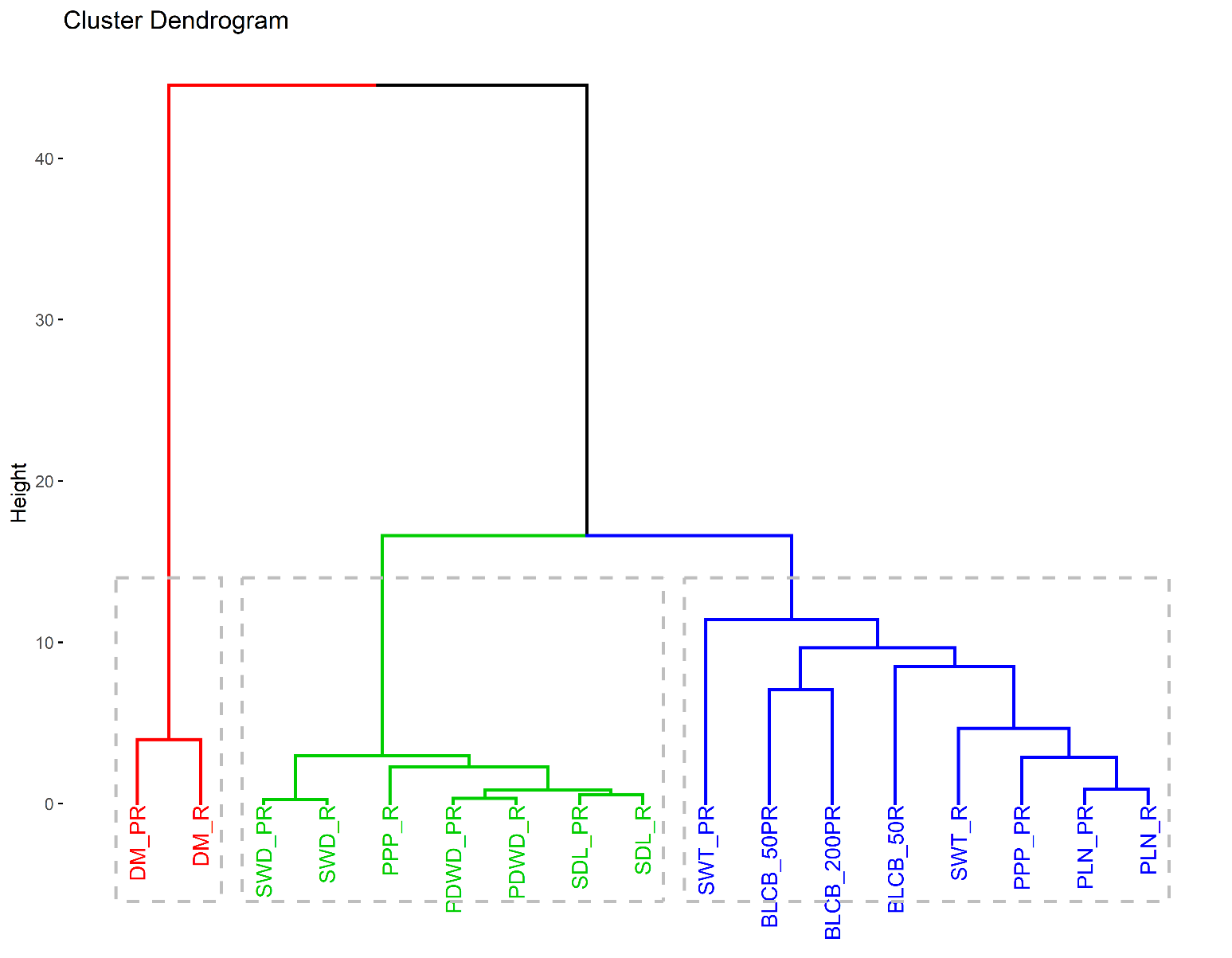
**Supplementary Figure S5: Hierarchical cluster analysis for blanchability and other agronomic traits** [DM (day to maturity), SWD (seed width), PPP (pods per plant), PDWD (pod width), SDL (seed length), SWT (seed weight), BLCB (Blanchability) (50 gm) (200 gm), PLN (plant length) (PR:post-rainy and R:rainy)] were grouped into three distinct clusters, are represented in different colors : **(1)** DM (PR and R); **(2)** SWD (PR and R),PPP (R), PDWD (PR and R), SDL (PR and R); **(3)** SWT (PR and R) (BLCB 50 (PR and R), 200 (PR), PPP (PR), PLN (PR and R)

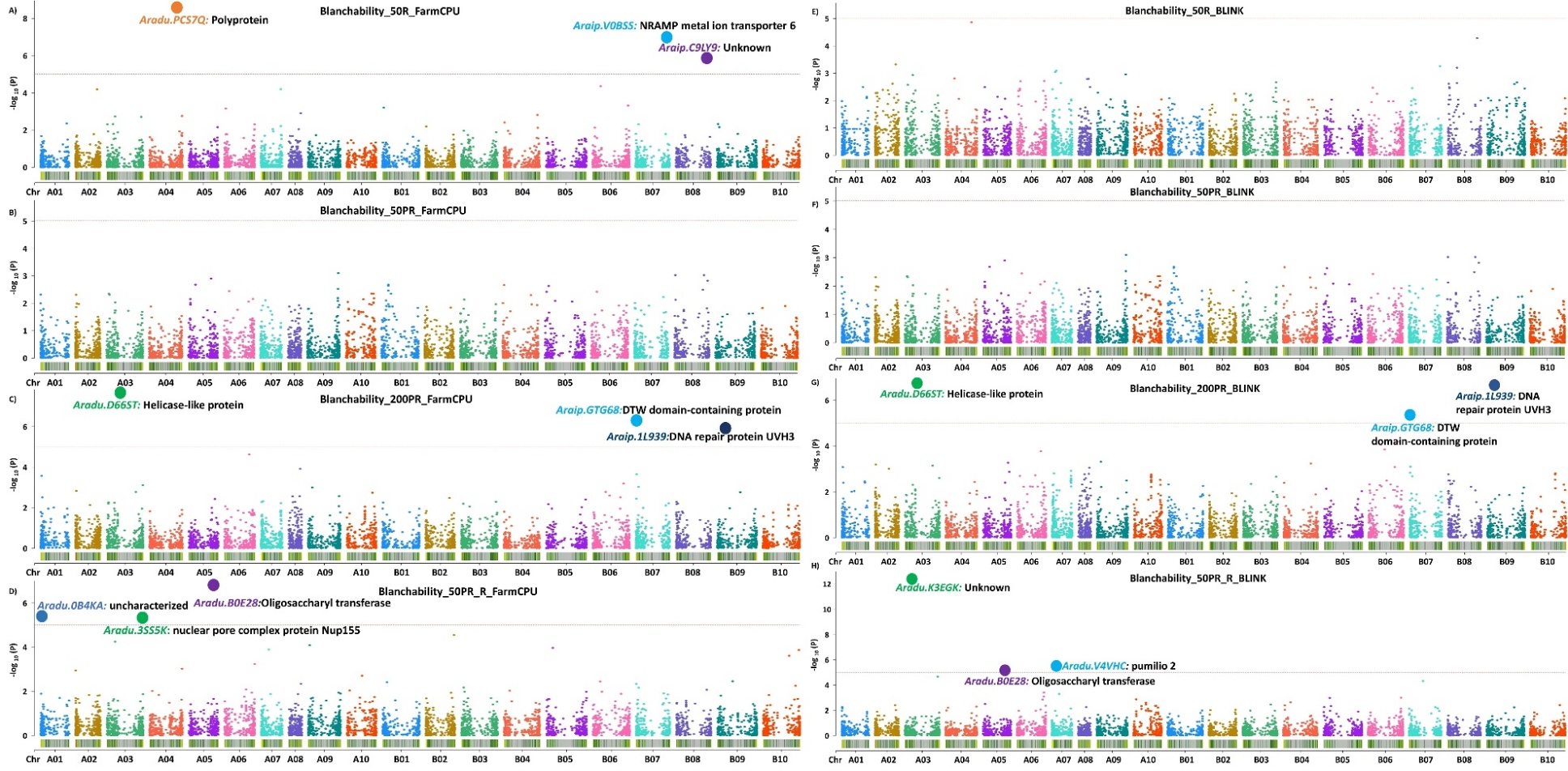

**Supplementary Figure S6:** **Manhattan plot representing the identified STAs associated with the blanchability on the basis A, B, C, D**: FarmCPU and **E, F, G, H**: BLINK model in GAPIT using 58K ‘Axiom_*Arachis*’ array data for Blanchability 50R, 50PR, 200PR and 50PR_R, respectively. The dots represent the significant STAs and the candidate genes.

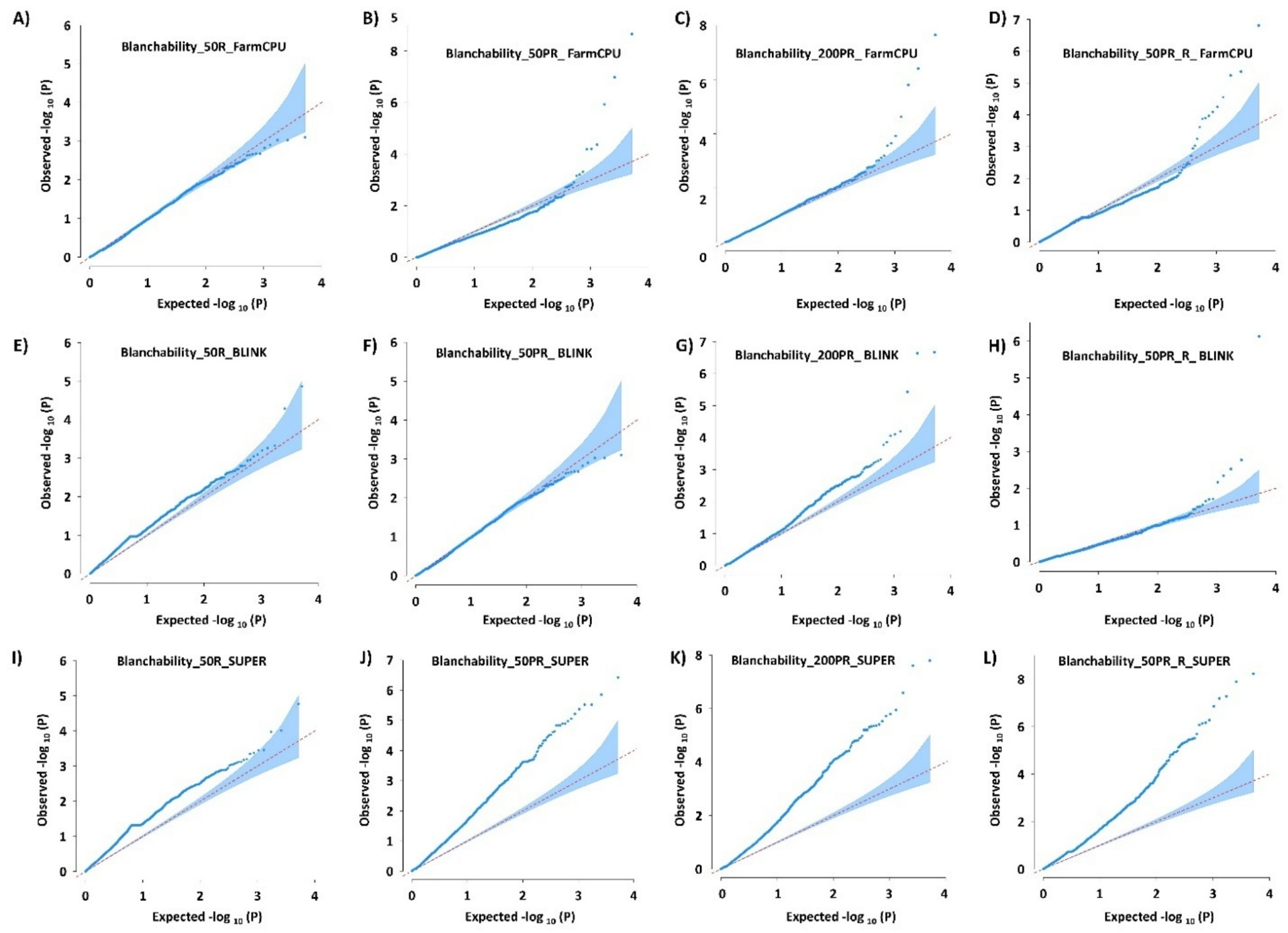

**Supplementary. Figure S7:** **Quantile-Quantile (Q-Q) plot representing the identified STAs associated with the blanchability on the basis A, B, C, D**: Q-Q plot representing the identified STAs associated with the blanchability on the basis FarmCPU and **E, F, G, H**: BLINK, and I, J, K, L: SUPER model in GAPIT, using 58K ‘Axiom_*Arachis*’ array data for Blanchability 50R, 50PR, 200PR and 50PR_R, respectively.

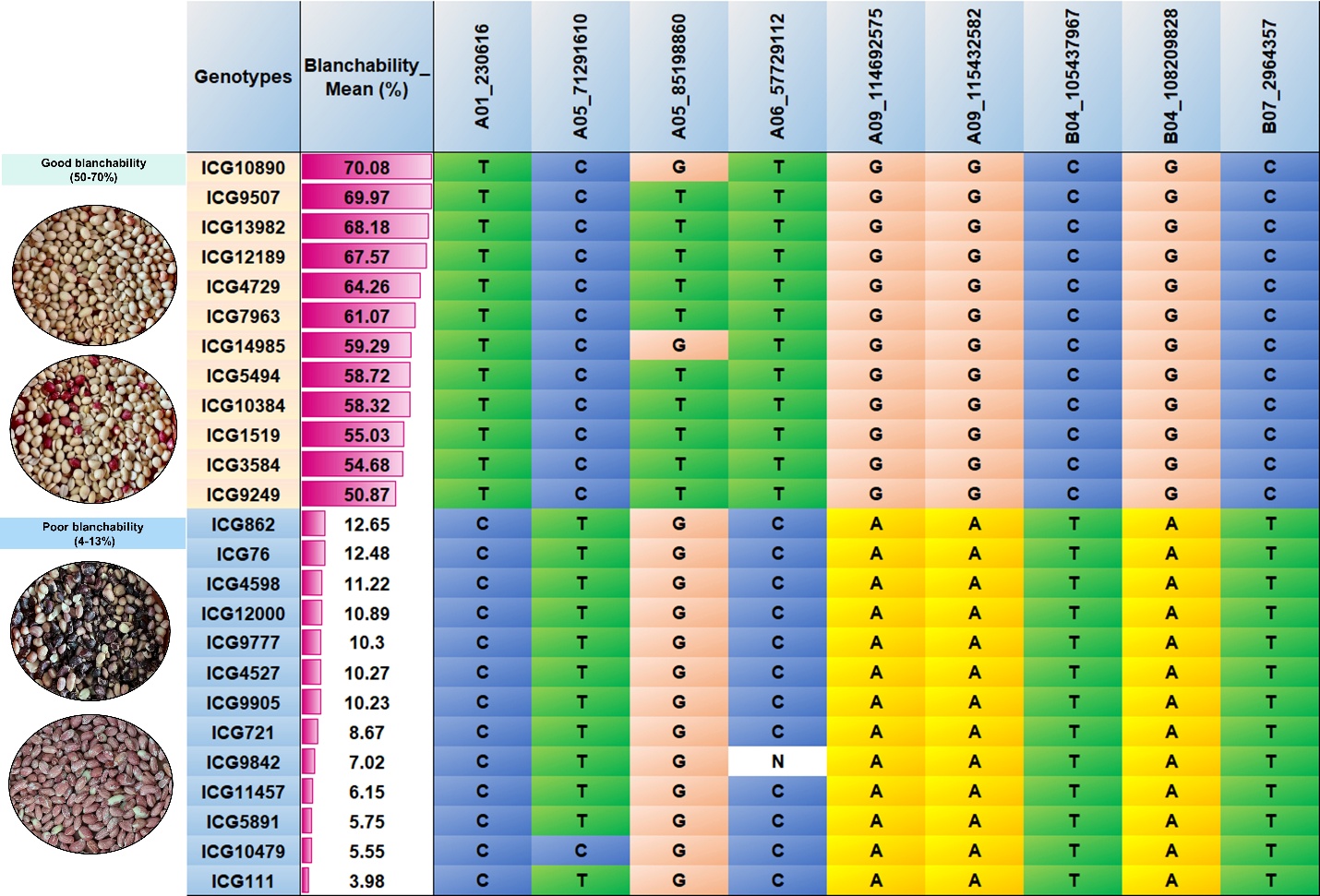

**Supplementary Figure S8: Nine stable signiﬁcant polymorphic SNPs for blanchability in minicore collection,** including; A01_230616, A05_71291610, A05_85198860, A06_57729112, A09_114692575, A09_115432582, B04_105437967, B04_108209828, B07_2964357

.
